# Prion Protein Deficiency Results in Synaptic, Neural Network and Behavioral Alterations

**DOI:** 10.64898/2026.04.07.716931

**Authors:** Anna Burato, Alessio di Clemente, Camilla Lodetti, Valentino Panico, Giulio Pistorio, Beatriz Eymi Pimentel Mizusaki, Beatrice Pastore, Marco Zattoni, Luigi Celauro, Lucia Zanetti, Lorenca Sadiraj, Eugenio Piasini, Michele Giugliano, Katja Reinhard, Giuseppe Legname

## Abstract

The cellular form of the prion protein (PrP^C^) is known for its involvement in the pathogenesis of prion diseases. Recent research implicates the physiological isoform of PrP in neuronal development, excitability, and synaptic plasticity, as well as in other biological processes. However, its precise function in the development and function of neurons remains poorly understood. Here, we investigated its role during different developmental stages, both *in vitro* and *in vivo*, using different PrP knock-out (KO) mouse lines (*Prnp^-/-^*). Prion protein KO neurons cultured on microelectrode arrays (MEAs) displayed altered network dynamics compared to wild type cultures, comprising reduced burst frequency, and abnormal spike patterns, indicative of impaired function of the synaptic circuitry. These functional alterations were associated with a reduced expression of key presynaptic and postsynaptic proteins, including elements of the SNARE complex and regulators of excitation-inhibition balance. Similar molecular changes were also confirmed in a second *Prnp^-/-^*model, suggesting that PrP^C^ is directly involved in these mechanisms regardless of genetic backgrounds. Alterations in neuronal networks were traceable into adulthood: *in vivo* recordings in adult *Prnp^-/-^* mice revealed increased neuronal responses to visual danger stimuli, which correlated with behaviorally increased fear responses to those stimuli. Together, our findings support a critical role for PrP^C^ in the maintenance of functional neuronal networks, from mature cortical neurons *in vitro* to behaviorally mature relevant circuits *in vivo*, beyond genomic background. These results indicate that PrP^C^ acts as a key regulator of synaptic function both in physiological and pathological conditions.

**Graphical Abstract:** 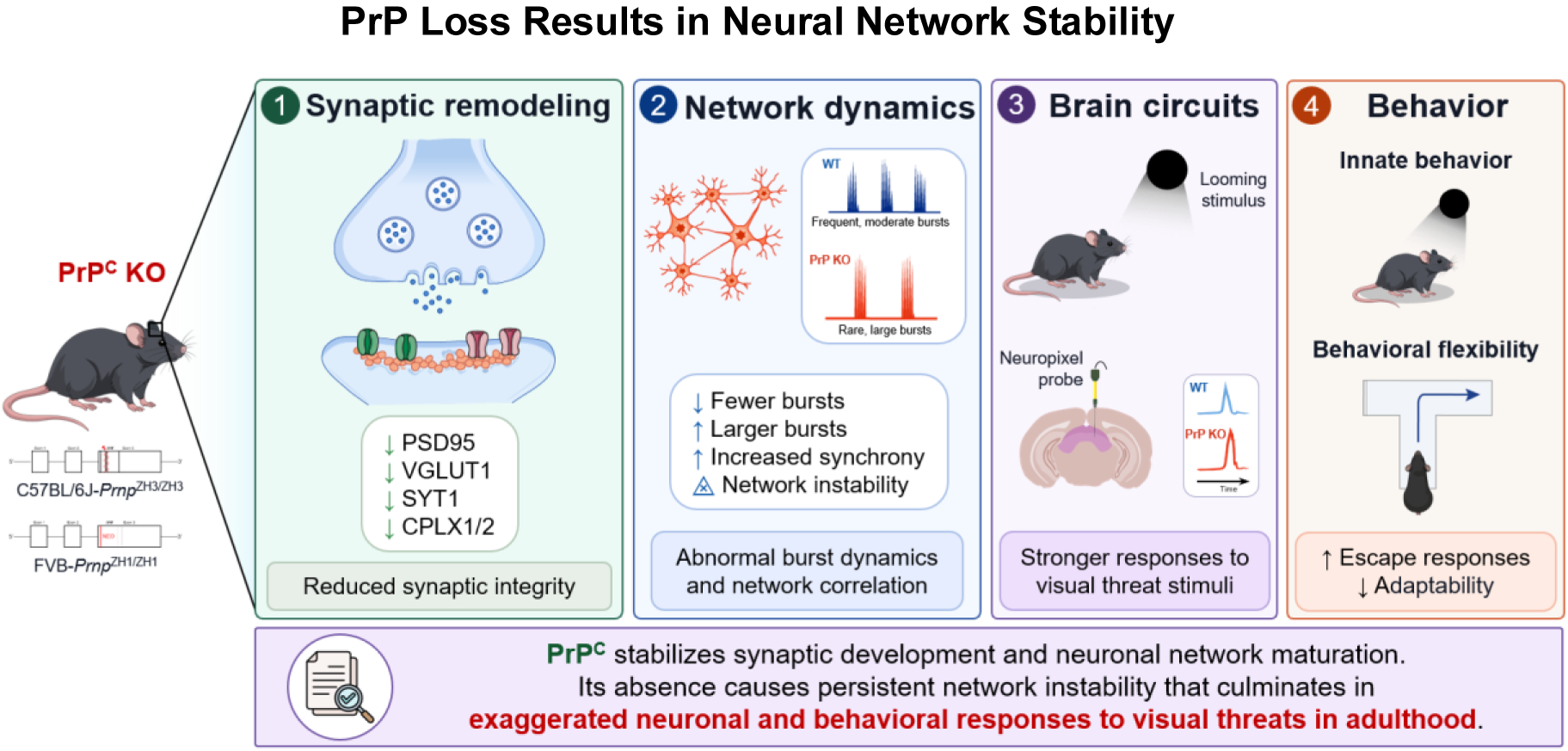

Prion protein deficiency is associated with altered synaptic protein expression and time-dependent differences in network burst dynamics, together with persistent alterations in adult neural activity and behavioral differences. Across two independent knockout mouse models, these findings indicate that chronic PrP loss is accompanied by measurable neuronal phenotypes that may inform the evaluation of PrP lowering strategies

## Introduction

The cellular prion protein (PrP^C^) is a ubiquitously expressed protein particularly abundant in brain tissue, including in neuronal and various non-neuronal cells such as astrocytes, oligodendrocytes, and microglia ^1–9^. Furthermore, its expression has been reported in immune lymphocytes and mast cells ^10,11^, and in many other peripheral body compartments such as heart, liver, intestine and kidney ^3,5,6,10^. This protein has been the center of research efforts, as its pathological conversion into a misfolded isoform, known as PrP scrapie (PrP^Sc^) ^12^, is the common denominator of prion diseases, a group of fatal neurodegenerative disorders. Prion diseases remain incurable, and the most promising strategy to date is targeting the physiological isoform, PrP^C^, since, due to the heterogeneity of prion strains, targeting the misfolded isoform directly is not feasible. However, a critical question remains: what are the physiological consequences of losing the cellular prion protein?

No strong recognizable phenotypes are associated with the lack of PrP^C^ ^13^, however, its widespread expression suggests that this protein is still involved in several physiological functions^5^. Phenotypic studies of *Prnp*-ablated mice suggest a role of PrP^C^ in neuritogenesis ^14–16^, cell signaling ^17–20^, and cell adhesion ^15,21,22^. Other studies reported an involvement in the response against stress ^23–25^, in circadian rhythms ^26^, and in sleep dysfunction ^27^. And finally, *Prnp*-ablated mice show abnormalities in neural stem differentiation in the central nervous system ^7,27–29^, in myelination in the peripheral nervous system^30^, and in synaptic plasticity ^31^.

The specifics of the physiological role of PrP^C^ remain enigmatic, largely due to the strong genetic confounders present in most *Prnp^-/-^* mouse lines. The two primary and most consistent lines used in recent functional researches are the Friend leukaemia virus B sensitive (FVB.*Prnp^0/0^*) mouse line, which is not fully co-isogenic and may retain flanking genetic material linked to the targeted *Prnp* locus, and the co-isogenic C57BL/6J-*Prnp^−/−^* mouse model (Zurich-3, ZH3), which was generated on a pure C57BL/6J background ^32^. Molecular and phenotypic studies on the new ZH3 model confirmed some of the previous data acquired with the FVB line, such as the role of PrP^C^ in myelin maintenance ^32^ and in regulating neuronal network excitability, formation, and connectivity, as well as in complex cognitive functions such as associative learning and anxiety-like behavior ^33^.

This suggests that in this scenario both lines are a valuable tool to study the implication of PrP^C^ in the structural and functional development of neurons and neuronal networks.

In the healthy brain, however, studies directly interrogating how the absence of PrP^C^ affects spontaneous neuronal activity, synaptic coordination, and the molecular machinery governing synaptic vesicle cycling are limited. This is particularly relevant considering therapeutic strategies that aim to reduce PrP^C^ expression to halt prion propagation. PrP^C^ is involved in multiple synaptic mechanisms. It supports synapse formation and stabilization ^34^, promotes neurite outgrowth and protects against oxidative stress-induced cytoskeletal disruption ^5^. By regulating calcium signaling, it ensures that vesicles are properly docked and primed for release via SNARE complex stabilization ^35^, and it supports synaptic homeostasis by buffering copper, regulating oxidative stress ^36,37^ and, through its interaction with copper, limiting the neurotoxic effects of NMDAR’s overactivation at glutamatergic synapses ^19^. Also, it regulates excitatory-inhibitory balance through NMDA and other receptor signaling ^35,38–40^. The interaction between the prion protein and the synaptic machinery is further confirmed by the synaptic degeneration found in prion-disease models, where the cellular form becomes less available to possibly regulate the pathway ^41–43^.

Since prion strains exhibit distinct pathological conformations, therapeutic strategies have taken an indirect approach, such as reducing PrP^C^ expression ^44,45^ or enhancing its degradation ^46,47^. PrP^C^ serves as the substrate for PrP^Sc^ propagation, and its removal effectively halts disease progression, as evidenced by resistance to prion diseases in PrP KO animals inoculated with prions ^48^. However, studies directly interrogating how the absence of PrP^C^ affects spontaneous neuronal activity, synaptic coordination, and the molecular machinery governing synaptic vesicle cycling are limited. Here, we investigated how PrP^C^ deficiency affects neuronal network maturation and function across developmental stages using both *in vitro* and *in vivo* models, and across distinct genetic backgrounds. The novelty of our study lies in the integration of synaptic molecular profiling, longitudinal MEA network analysis, adult *in vivo* electrophysiology, and behavioral assays within the same experimental framework. This multilevel approach allowed us to link PrP^C^ deficiency to convergent alterations in synaptic organization, network dynamics, and stimulus-evoked behavioral responses, while distinguishing shared effects of PrP loss from model-specific features. We identify PrP^C^ as a key regulator of neuronal network maturation and stability, whose loss is linked to persistent instability, rarer and abnormal population events, increased pairwise rate correlation and impaired plasticity and behavior into adulthood. These findings establish disrupted network control as a related consequence of PrP^C^ deficiency and a critical consideration for PrP^C^ targeting therapies.

## Results

### PrP^C^ Deletion Impairs Synaptic Network Activity in Mature Cortical Cultures

To characterize network activity across *in vitro* maturation, cortical cultures from either wild-type (hereafter wtFVB) or FVB.*Prnp*^0/0^ (hereafter koFVB) mice were plated on 120-electrode multielectrode arrays (MEAs) (see Methods). Absence of PrP in koFVB cultures was confirmed by PrP immunostaining (Figure S1A), genotyping (Figure S1B), real-time qPCR and western blotting (Figure S1C). Although the cultures showed structural differences at earlier stages (Figure S1D–E), neuronal density at maturation did not differ between wtFVB and koFVB cultures, suggesting that cell survival was not a confounder (Figure S1G–H). By contrast, quantification of excitatory synapses (VGLUT1/PSD-95 puncta) at maturation revealed reduced excitatory synapse density in koFVB cultures relative to wtFVB (Figure S2).

Network activity was recorded at three developmental time points: DIV10, DIV17, and DIV23. These time points were chosen to capture three stages of *in vitro* network maturation: emergence of network activity (DIV10), functional refinement (DIV17), and mature network activity (DIV23) (van Pelt, Vajda et al. 2005). Spontaneous activity was recorded for 30 min per MEA at each time point. Representative raster plots and spike-time histograms (STHs) illustrate qualitative differences in activity patterns at the chosen developmental stages (Figure 1A–B). Despite these differences, the total spike count per MEA over the 30-min recording was comparable between wtFVB and koFVB cultures at DIV10, DIV17, and DIV23 (Figure 1C). By contrast, the occurrence of network-wide bursts (“network bursts”) (van Pelt, Vajda et al. 2005; Wagenaar, Pine et al. 2006) was significantly reduced in koFVB cultures at maturation, as reflected by longer inter-burst intervals (IBIs) at DIV23 (median of MEA medians: wtFVB, 3395 ms; koFVB, 9518 ms; adjusted p = 0.0260; *r_rb_ = −0.635; Figure 1D)*.

**Figure 1:**
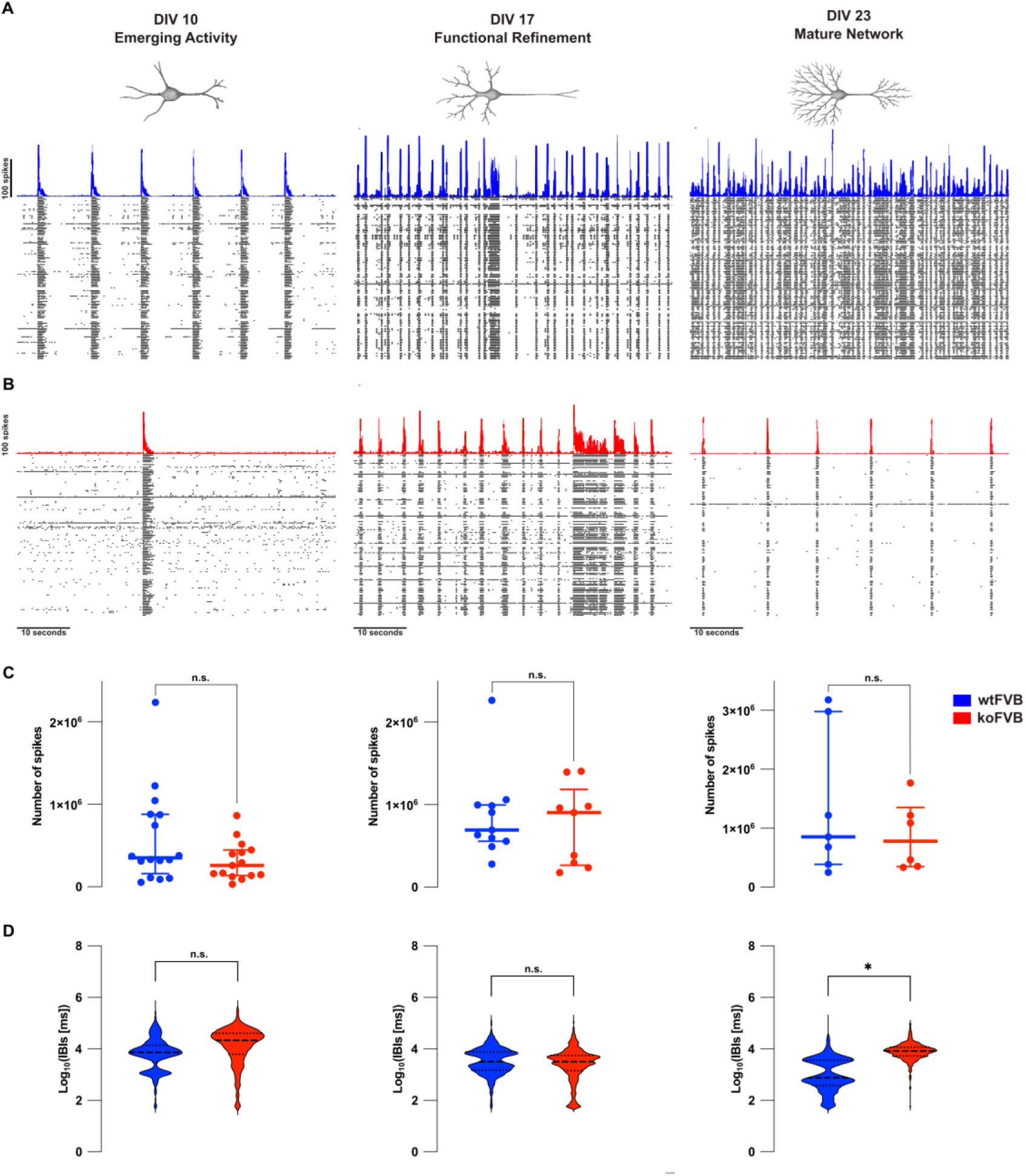
Reduced network-burst occurrence at maturation in koFVB cortical cultures. (A–B) Representative multielectrode-array (MEA) recordings from cortical cultures at DIV10, DIV17, and DIV23 showing spike rasters (black) with the corresponding spike-time histograms (STHs; 5-ms bins; wtFVB, blue in A; koFVB, red in B). Scale bars as indicated. (C) Total spike count per MEA during 30-min spontaneous recordings at each developmental stage. Each dot represents one MEA; horizontal lines indicate median with interquartile range (IQR). DIV10: Mann–Whitney U test, two-tailed, p = 0.2641 (wtFVB n = 16 MEAs, koFVB n = 15 MEAs). DIV17: Mann–Whitney U test, two-tailed, p = 0.6027 (wtFVB n = 11, koFVB n = 9). DIV23: unpaired t test with Welch’s correction, two-tailed, p = 0.3654 (wtFVB n = 7, koFVB n = 6). (D) Inter-burst interval (IBI) distributions for network-wide bursts at DIV10, DIV17, and DIV23. Violin plots display log10-transformed IBI values (ms) for visualization only; statistics were performed on raw, untransformed IBI values. DIV10: clustered Wilcoxon, adjusted p = 0.4611 (wtFVB n = 16 MEAs, 1583 IBIs; koFVB n = 15 MEAs, 803 IBIs; median of MEA medians = 14,671 vs 20,230 ms; r_rb_ = −0.140). DIV17: clustered Wilcoxon, adjusted p = 0.9968 (wtFVB n = 11 MEAs, 2724 IBIs; koFVB n = 8 MEAs, 2178 IBIs; median of MEA medians = 6553 vs 5955 ms; r_rb_ = −0.001). DIV23: clustered Wilcoxon, adjusted p = 0.0260 (wtFVB n = 6 MEAs, 2454 IBIs; koFVB n = 6 MEAs, 858 IBIs; median of MEA medians = 3395 vs 9518 ms; r_rb_ = −0.635). Effect size is reported as the cluster-weighted rank-biserial correlation (r_rb_). Data from 3 independent culture preparations. *p ≤ 0.05; **p ≤ 0.01; ***p ≤ 0.001; ****p ≤ 0.0001; n.s., not significant.

Average spike-time histograms aligned to burst onset (Figure 2A) showed qualitative differences in burst profiles between wtFVB and koFVB cultures. These differences were particularly evident at maturation. Quantitative analysis showed a tendency toward lower burst amplitudes in koFVB cultures at DIV10. The same pattern was more pronounced at DIV17, with stronger statistical support (adjusted p = 0.0500; r_rb_ = 0.436). By DIV23, the direction of this difference was reversed, although no significant difference was detected. Burst onset slope and duration did not differ significantly between wtFVB and koFVB cultures at any developmental stage. To assess network synchrony, we computed pairwise rate correlations across electrodes and summarized the distribution of significant cross-correlation indices for each condition (Figure 2E). No significant genotype differences were detected at DIV10 or DIV17. By contrast, at DIV23, koFVB cultures exhibited higher cross-correlation indices overall (adjusted p = 0.0476; r_rb_ =−0.656; Figure 2E).

**Figure 2:**
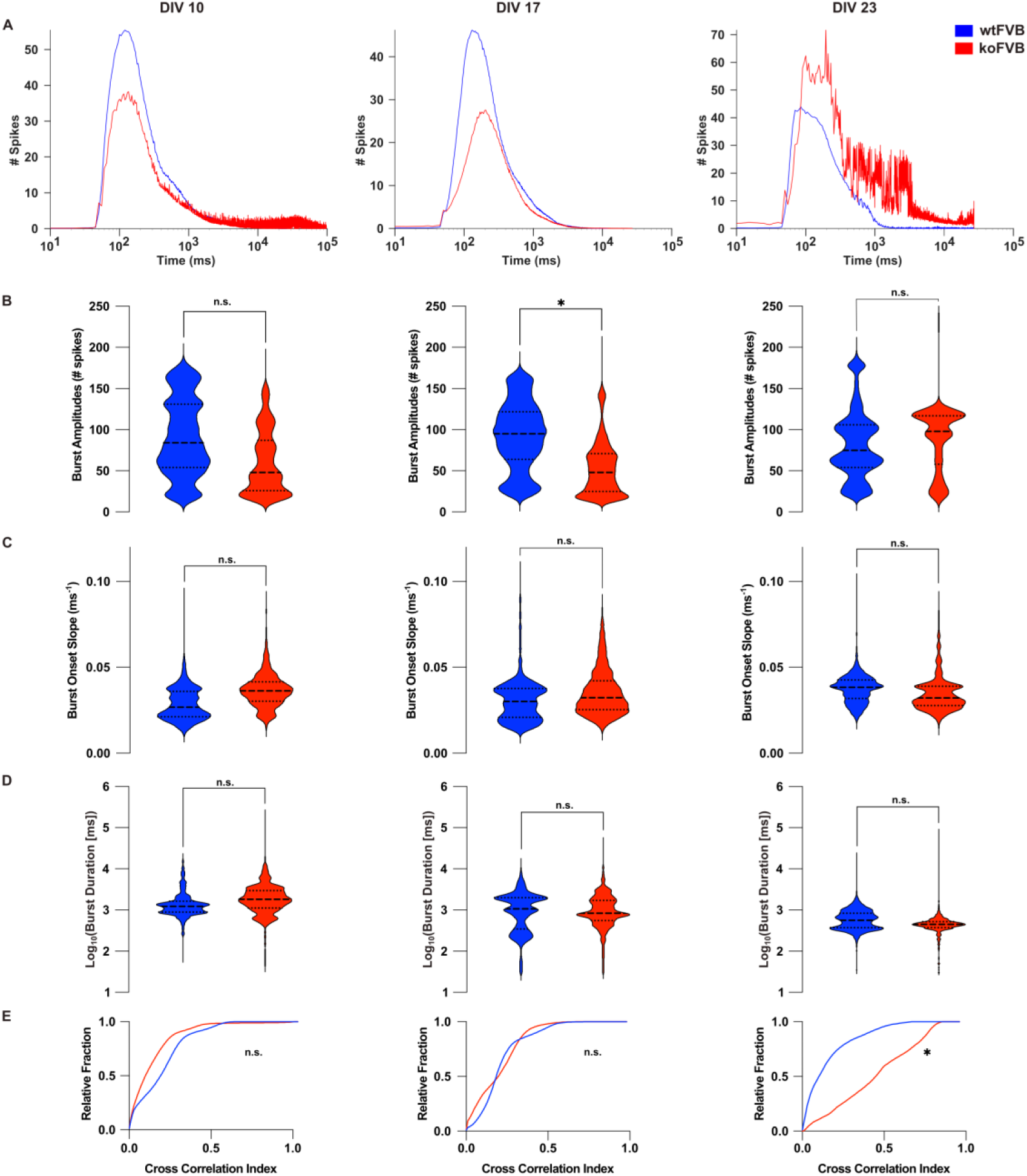
Burst profiles and rate correlation differ between wtFVB and koFVB cortical cultures. (A) Average spike-time histograms (STHs; 5-ms bins) aligned to network-burst onset at DIV10, DIV17, and DIV23. Traces show the mean across all detected network bursts pooled across MEAs for each genotype and time point (wtFVB, blue; koFVB, red). (B–D) Burst amplitude (B; spike count in the peak bin), burst onset slope (C; maximal rising slope at burst onset, ms⁻¹), and burst duration (D; ms) for individual network bursts pooled across MEAs for each condition and time point. Violin plots show distributions; horizontal lines indicate median and interquartile range (IQR). Burst duration values are log10-transformed for visualization only; statistics were performed on raw, untransformed values. (B) DIV10: clustered Wilcoxon, adjusted p = 0.3011 (wtFVB n = 16 MEAs, 1599 bursts; koFVB n = 15 MEAs, 818 bursts; median of MEA medians = 87.25 vs 51 spikes; r_rb_ = 0.265). DIV17: clustered Wilcoxon, adjusted p = 0.0500 (wtFVB n = 11 MEAs, 2735 bursts; koFVB n = 8 MEAs, 2186 bursts; median of MEA medians = 83 vs 51.5 spikes; r_rb_ = 0.436). DIV23: clustered Wilcoxon, adjusted p = 0.4026 (wtFVB n = 6 MEAs, 2460 bursts; koFVB n = 6 MEAs, 864 bursts; median of MEA medians = 55.5 vs 106.5 spikes; r_rb_ = −0.267). (C) DIV10: clustered Wilcoxon, adjusted p = 0.5953 (wtFVB n = 16 MEAs, 1599 bursts; koFVB n = 15 MEAs, 815 bursts; median of MEA medians = 0.03473 vs 0.03659 ms⁻¹; r_rb_ = −0.164). DIV17: clustered Wilcoxon, adjusted p = 0.5953 (wtFVB n = 11 MEAs, 2729 bursts; koFVB n = 8 MEAs, 2183 bursts; median of MEA medians = 0.02609 vs 0.03135 ms⁻¹; r_rb_ = −0.209). DIV23: clustered Wilcoxon, adjusted p = 0.4416 (wtFVB n = 6 MEAs, 2409 bursts; koFVB n = 6 MEAs, 864 bursts; median of MEA medians = 0.04182 vs 0.03472 ms⁻¹; r_rb_ = 0.378). (D) DIV10: clustered Wilcoxon, adjusted p = 1.000 (wtFVB n = 16 MEAs, 1599 bursts; koFVB n = 15 MEAs, 818 bursts; median of MEA medians = 1332.5 vs 1775 ms; r_rb_ = −0.111). DIV17: clustered Wilcoxon, adjusted p = 1.000 (wtFVB n = 11 MEAs, 2735 bursts; koFVB n = 8 MEAs, 2186 bursts; median of MEA medians = 1810 vs 1230 ms; r_rb_ = 0.182). DIV23: clustered Wilcoxon, adjusted p = 1.000 (wtFVB n = 6 MEAs, 2460 bursts; koFVB n = 6 MEAs, 864 *bursts*; median of MEA medians = 718.75 vs 461.25 ms; r_rb_ = 0.032). (E) Cumulative distributions of cross-correlation indices computed from pairwise rate correlations across electrodes; only electrode pairs significant against a null distribution (see Methods) are shown (wtFVB, blue; koFVB, red). DIV10: clustered Wilcoxon, adjusted p = 0.2742 (wtFVB n = 16 MEAs, 90,853 electrode pairs; koFVB n = 15 MEAs, 87,720 pairs; median of MEA medians = 0.2050 vs 0.09364; r_rb_ = 0.251). DIV17: clustered Wilcoxon, adjusted p = 0.4224 (wtFVB n = 10 MEAs, 64,759 pairs; koFVB n = 9 MEAs, 49,573 pairs; median of MEA medians = 0.1948 vs 0.1877; r_rb_ = 0.158). DIV23: clustered Wilcoxon, adjusted p = 0.04762 (wtFVB n = 5 MEAs, 28,862 pairs; koFVB n = 5 MEAs, 31,418 pairs; median of MEA medians = 0.09097 vs 0.4480; r_rb_ = −0.656). Effect size is reported as the cluster-weighted rank-biserial correlation (r_rb_). Data from 3 independent culture preparations. *p ≤ 0.05; **p ≤ 0.01; ***p ≤ 0.001; ****p ≤ 0.0001; n.s., not significant.

### Synaptic pathways are disrupted in PrP^C^ deficient neurons at late maturation stage

To identify the molecular pathways accompanying the functional phenotype observed in mature koFVB neuronal networks, we performed RNA-sequencing on DIV23 primary cortical cultures (Figure S3). Differential expression analysis highlighted consistent alterations in genes related to cytoskeletal organization, synaptic efficacy, and neurotransmission dynamics (Table S2), suggesting that PrP^C^ loss impacts molecular elements required for maintaining stable synaptic connectivity and efficient transmission once networks reach maturity.

Guided by these transcriptomic signatures, we quantified key pre- and postsynaptic proteins in wtFVB and koFVB neurons. Immunoblotting confirmed effective PrP^C^ ablation and revealed coordinated changes in synaptic markers in koFVB primary cultures across development (Figure 3B, D). At the level of the postsynaptic compartment, PSD95 showed a progressive reduction in koFVB cultures, with the clearest deficit detected at later stages when synapses are stabilized (DIV23) (Figure 3B, D); similarly, we found a selective decrease in the glutamatergic presynaptic marker VGLUT1 (Figure 3B, D). Immunofluorescence confirmed that active glutamatergic synapses are less abundant in koFVB at the maturation stage (Figure S2).

**Figure 3:**
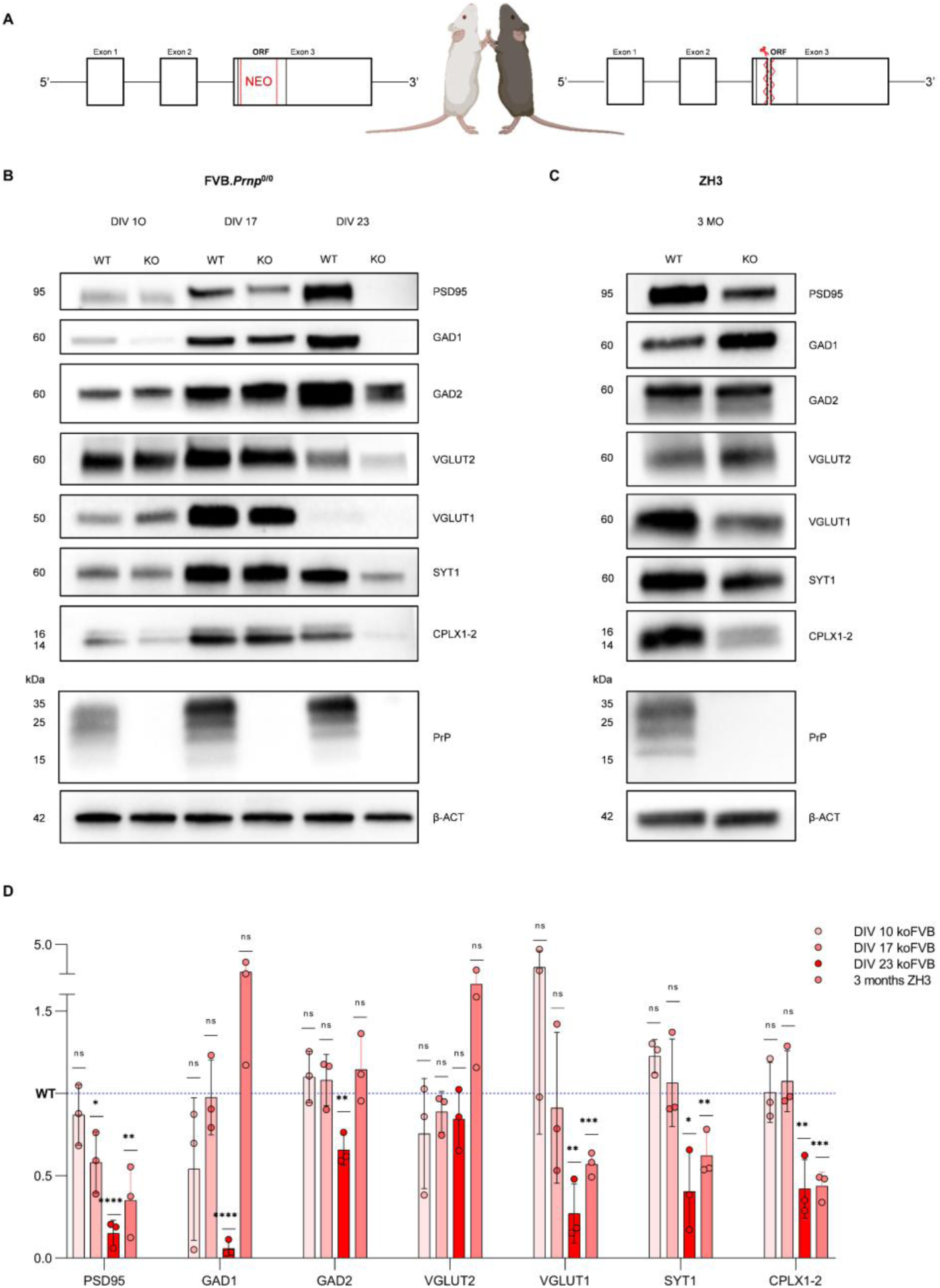
Synaptic protein changes across maturation development. (A) Schematic of the strategies used to generate the FVB (left) and the ZH3 KO lines (right). (B) Representative immunoblots from FVB cortical primary cultures at DIV10, DIV17, and DIV23 probed for synaptic and neurotransmission related markers: PSD95 (DIV10 p = 0.2831, DIV17 p = 0.0176, DIV23 p<0.0001), GAD1 (DIV10 p = 0.1398, DIV17 p = 0.9657, DIV23 p<0.0001), GAD2 (DIV10 p = 0.3256, DIV17 p = 0.4173, DIV23 p = 0.0031), VGLUT2 (DIV10 p = 0.2745, DIV17 p = 0.1920, DIV23 p = 0.2209), VGLUT1 (DIV10 p = 0.2100, DIV17 p = 0.7570, DIV23 p = 0.0022), SYT1 (DIV10 p = 0.0178, DIV17 p = 0.6953, DIV23 p = 0.0132), CPLX1–2 (DIV10 p = 0.9571, DIV17 p = 0.5264, DIV23 p = 0.0050), together with PrP to confirm knockout and β-actin as loading control. Molecular weight markers (kDa) are indicated. (C) Representative immunoblots from adult ZH3 cortex (3 months) probed for the same panel of proteins: PSD95 (p = 0.0065), VGLUT1 (p = 0.0005), VGLUT2 (p = 0.1018), GAD1 (p = 0.1436), GAD2 (p = 0.2938), SYT1 (p = 0.0092), CPLX1-2 (p = 0.0003). (D) Densitometric quantification of protein levels normalized to β-actin and expressed relative to WT (dashed line). Points represent individual biological replicates; bars show mean ± variability as displayed. Statistical significance was assessed by unpaired two-tailed t-test for WT vs KO within each condition/time point. *p ≤ 0.05. **p ≤ 0.01. ***p ≤ 0.001. ****p ≤ 0.0001, n.s. not significant. Full uncropped blots and relative actins are available in Table S4.

In parallel, within the presynaptic compartment, multiple proteins involved in vesicle release and synaptic transmission were reduced in koFVB neurons, including SYT1 and CPLX1–2. A similar reduction was observed for GAD1 and GAD2, consistent with changes in inhibitory-related molecular programs. Together with the changes in glutamatergic markers, these findings suggest a broad disruption of synaptic signaling (Figure 3B, D). In contrast, VGLUT2 expression was comparatively preserved across timepoints, indicating that PrP^C^ loss does not uniformly suppress neurotransmitter-related proteins but preferentially affects a subset of synaptic components (Figure 3B, D). In summary, we found a late emerging synaptic maintenance effect rather than an early developmental failure in PrP KO.

### Conserved glutamatergic and synaptic-marker changes are observed across KO strategies and in adult ZH3 cortex

We tested whether the molecular and electrophysiological alterations were specific to the FVB model or a general feature of prion protein deficiency. We hence performed RNA-sequencing on ZH3 derived primary cortical neurons cultured under the same conditions used for FVB cultures (DIV23), to separate background effects from the biological readout of PrP^C^ loss (Figure S4). In contrast to the broader transcriptomic dysregulation detected in koFVB neurons, ZH3 cultures showed no comparable alterations on the RNA level, consistent with possible additional transcriptional perturbations related to the targeting strategy and/or effects on neighboring genomic regions in the FVB line ^32^. We therefore used the ZH3 model as a genetically cleaner system to test whether the *in vitro* phenotype observed in koFVB cultures was truly attributable to PrP deficiency. Specifically, we repeated the electrophysiological experiments in ZH3 derived primary cultures under identical conditions. ZH3 cultures showed qualitatively similar alterations to those observed in koFVB cultures, including a tendency toward longer IBIs, although these differences did not reach statistical significance in the small ZH3 cohort (Figure S5) and a larger cohort would be required to strengthen the statistical support. Given the cleaner genetic background and comparable molecular and cellular changes, we used ZH3 mice for further experiments.

We next tested whether the protein level changes identified in mature cultures are preserved by profiling adult cortex from 3-month-old WT and ZH3 KO mice (hereafter ZH3) (Figure 3C, D). Notably, PSD95 and VGLUT1 were significantly reduced in ZH3 cortex (Figure 3C, D), mirroring the most consistent glutamatergic deficits observed in mature koFVB cultures, and supporting the conclusion that PrP^C^ loss impacts glutamatergic synaptic integrity across genetic contexts and both during *in vitro* and *in vivo* maturation. SYT1 and CPLX1-2 were also reduced in adult ZH3 tissue, identifying presynaptic release-associated proteins as an additional convergent molecular phenotype (Figure 3C, D). Together, these cross-model results indicate that while some markers may show context-dependent modulation, reductions in glutamatergic synaptic scaffolding and presynaptic release-associated proteins represent conserved molecular correlates of PrP^C^ deficiency.

### Differences in spontaneous neural activity also *in vivo*

Based on the strong differences in mature KO networks, we tested whether effects of prion protein deficiency on neural network development translate into physiological differences in the adult brain. To address this, we performed *in vivo* silicon probe recordings of the retrosplenial cortex, superior colliculus and periaqueductal gray of head-fixed ZH3 and WT mice that could spontaneously move on a floating platform (Figure 4A). Similar as in our *in vitro* recordings, we found longer bursts produced by ZH3 neurons compared to WT neurons (Figure 4B) and a brain area specific trend towards fewer bursts in ZH3 neurons (cortex - WT: 0.0094 bursts/s (median), ZH3: 0.0057 bursts/s, p = 0.442; periaqueductal gray - WT: 0.0009 bursts/s, ZH3: 0.0006 bursts/s, p = 0.034) (Figure 4C). In addition, we found an overall higher correlation between the activity of simultaneously recorded cortical neurons in ZH3 (Figure 4D). This suggests that network alterations due to the lack of PrP^C^ are traceable into adulthood.

**Figure 4:**
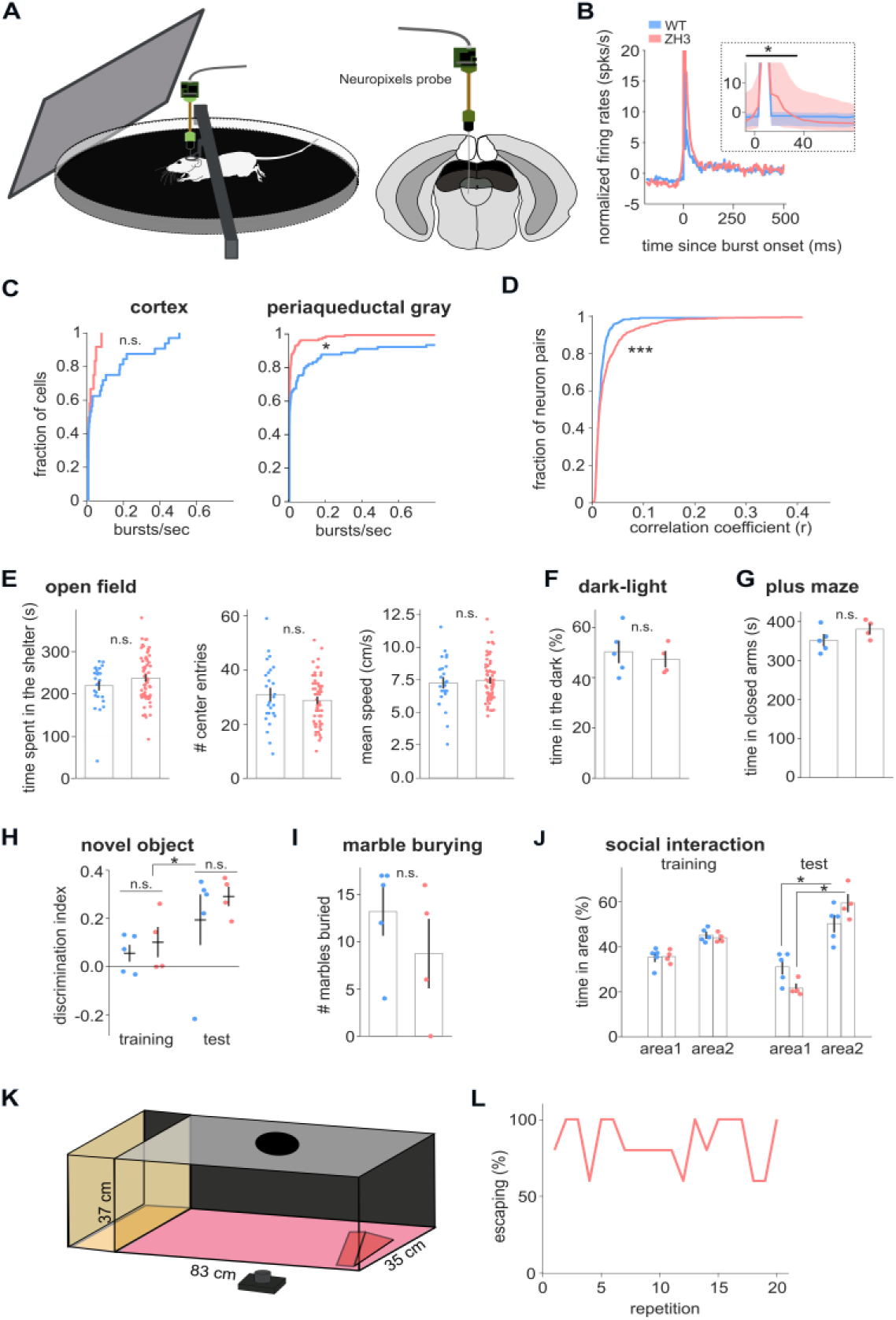
Effect of prion protein deficiency on in vivo neural and behavioral state in ZH3 mice. (A) Head-fixed setup with floating platform and Neuropixels probe (left). Recording sites are shown in a coronal slice. The probe passed through the retrosplenial cortex (white), the superficial (black) and deep (dark gray) layers of the superior *colliculus*, and the dorsal periaqueductal gray (lighter gray). (B) Mean normalized firing rate during bursts in WT (blue, n=6 mice) and ZH3 mice (red, n=5 mice). Inset shows median, 25% and 75% quantiles for -2 to 80 ms before/after burst onset. The bar on top indicates time bins for which WT and ZH3 are significantly different. (C) Burst rate per second across cortical (WT: n=32 cells, ZH3: n=12 cells; p = 0.442) and periaqueductal gray cells (WT: n=89, ZH3: n=129; p = 0.034). Statistical comparisons were performed using Wilcoxon-Ranksum Test. (D) Correlations between all pairs of recorded cells in the cortex. (E) Open-field assay (WT: n=26 trials from n=8 mice; ZH3: n=65 trials from n=14 mice). Time spent in the shelter (left), number of center entries (middle) and mean speed during 15 min of exploration (right) are shown. Bar plots indicate means, vertical lines indicate S.E.M. Shelter time WT: 221±10 s (mean ± sem), ZH3: 239±7 s p = 0.165; center entries WT: 30.9±1.1 entries, ZH3: 28.8±1.1 entries, p = 0.340; mean speed WT: 7.5±0.2 cm/s, ZH3: 7.3±0.4 cm/s p = 0.622. (F-J) Other behavioral assays (WT: n=5 mice, KOZH3: n=4 mice). (F) Time spent in the dark part of the dark-light chamber. WT: 50.3±4.2 s, ZH3: 47.3±3.0 s, p = 0.600. (G) Time spent in the closed arms of an elevated plus maze. WT: 351±14 s, ZH3: 380±12 s, p = 0.156). (H) Novel-object assay. A discrimination index for two identical objects (training) or one familiar and one novel object (test) was calculated. Horizontal lines indicate mean. Training discrimination index WT: 0.055±0.035, ZH3: 0.102±0.063, test discrimination index WT: 0.194±0.105, ZH3: 0.291±0.041; WT-ZH3 (test) p = 0.501, WT-ZH3 (training) p = 0.651. ANOVA was used *to* test for interactions between sessions and genotypes. Session p = 0.038, genotype p = 0.330, session*genotype p = 0.728. (I) Marble burying assay. WT: 13.2±2.5 marbles, ZH3: 8.8±3.6 marbles, p = 0.299. (J) Social interaction assay. On the test day, area2 contained a conspecific of the same sex. Training WT: 35.5±1.9% in area1, 45.0±1.3% in area2, ZH3: 35.7±1.4% in area1, 43.9±1.0% in area2; Test: WT: 30.9±3.0% in area1, 50.3±3.6% in area2 (with conspecific), ZH3: 21.5±1.7% in area1, 59.4±3.6% in area2; p (WT area1 vs. area2) < 0.001; p (ZH3 area1 vs. area2 = 0.001; p (WT vs. ZH3 area2) = 0.524. (K) Setup for looming stimuli. (L) Percent of escaping KO ZH3 mice during 20 repetitions of a looming stimulus (n=5). All vertical lines indicate standard deviation from the mean. All statistics were performed using DABEST permutation tests with Bonferroni correction where applicable, unless otherwise indicated. *p ≤ 0.05. **p ≤ 0.01. ***p ≤ 0.001. ****p ≤ 0.0001, n.s. not significant.

### Behavioral flexibility but not states are affected by PrP KO

To understand how general neural network alterations in the adult mouse may be linked to behavioral changes we performed a set of standardized assays. First, we aimed to test whether PrP KO affects behavioral states. We found no significant difference in parameters related to anxiety levels between ZH3 and WT mice when tested in an open-field assay (Figure 4E, Figure S6) in a dark-light chamber (Figure 4F), an elevated plus maze (Figure 4G), or when partially restricted during head-fixation on a floating platform (session 1 in Figure S6). Furthermore, we found no effect of PrP KO on exploration of inanimate objects where both genotypes explored novel objects equally more than familiar ones (Figure 4H). Similarly, no differences were found for marble burying (Figure 4I) or social interactions (Figure 4J). PrP^C^ deficiency hence does not appear to affect behavioral states such as anxiety, sociability and curiosity of adult mice.

To test the effect of PrP KO on behavioral flexibility, ZH3 mice were placed in an arena and exposed to many repetitions of a visual looming stimulus mimicking an overhead attacking predator (Figure 4K). While this stimulus initially evokes strong aversive behaviors, repeating it multiple times has been shown to induce quick neuronal and behavioral habituation ^49,50^. However, ZH3 mice did not habituate and kept escaping for up to 20 repetitions at consistent or even shorter latencies compared to the first presentation (Figure 4L; Figure S6C). These findings suggest a decrease in behavioral flexibility of ZH3 mice (see also Figure S6D-F).

### Behavioral and neuronal responses to danger stimuli are stronger in ZH3 mice

Finally, we asked whether PrP^C^ deficiency not only alters learned but also innate evoked behaviors. To do so, ZH3 and WT animals were placed in an open arena with a shelter where, after an acclimation phase, a black looming stimulus was presented (Figure 4K, 5A). ZH3 mice escaped earlier to the stimulus than wildtype littermates (latency to escape WT: 1.56±0.37 s, ZH3: 0.94±0.15s, p = 0.058 Permutation Test; Figure 5B; Figure S6) and showed a tendency to escape more often (67% vs 50%; generalized linear mixed-effect model, genotype effect β = 0.67, SE = 1.04, p = 0.52). This difference was even more pronounced in response to a dimming disk, which is considered a “control” stimulus without ethological meaning^51^. The dimming disk produced very few reactions in our WT animals, but elicited escape in significantly more ZH3 mice (63% vs. 27%; β = 1.62, SE = 0.78, p = 0.04) (Figure 5C-D). These findings suggest that PrP^C^ deficiency not only affects learning-related processes but also enhances innate evoked behaviors, in particular to low-danger stimuli.

**Figure 5:**
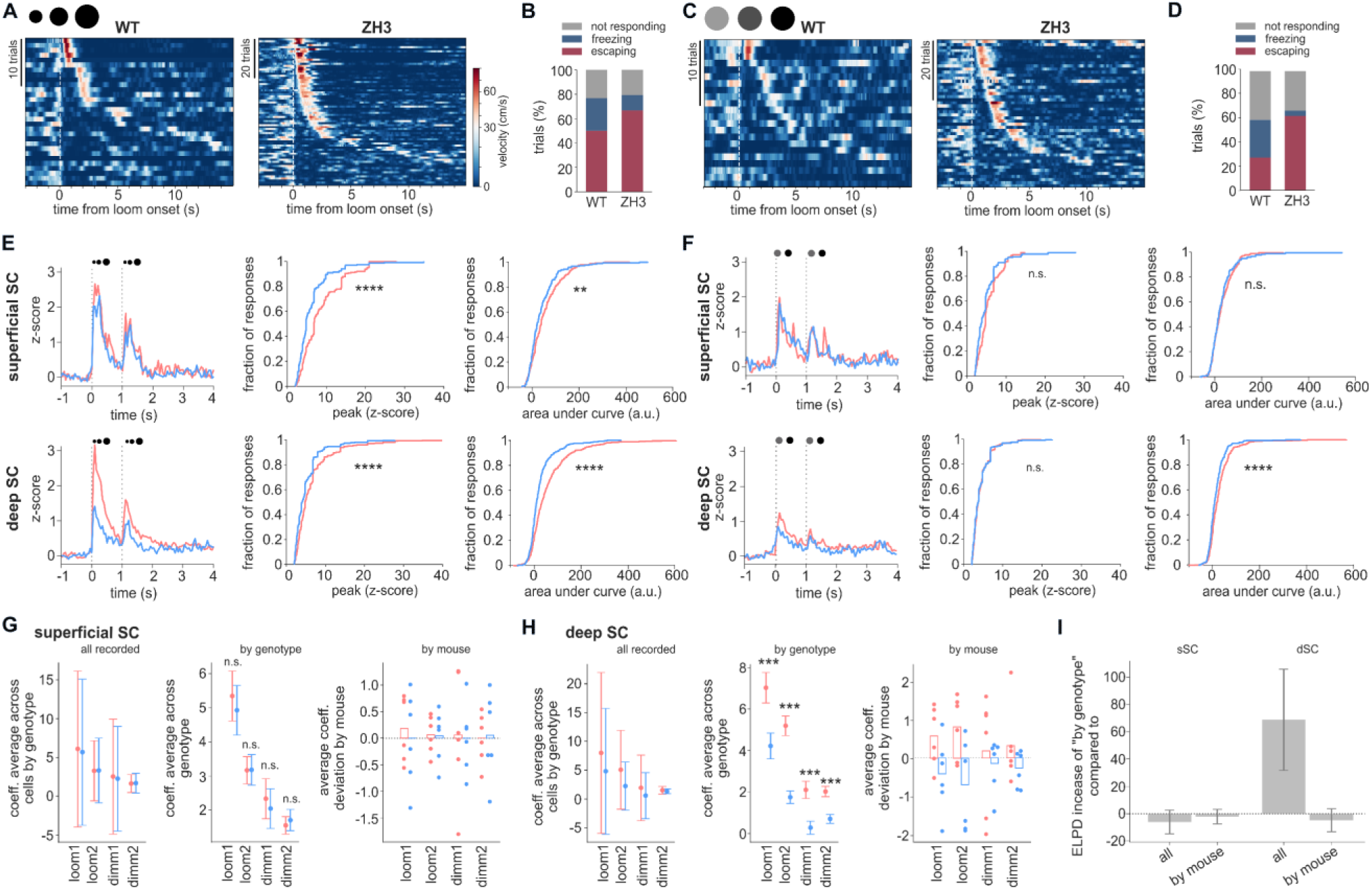
Aversive behavior and underlying neuronal activity is increased in ZH3 mice. (A) Heatmap of running velocity during the presentation of black looming stimuli. WT: n=30 trials from n=8 animals; ZH3: n=72 trials from n=14 animals. (B) Percentage of escape and freezing behavior during looming stimulus. (C-D) Same as A-B but for a dimming stimulus. WT: n=22 trials from n=8 animals; ZH3: n=48 trials from n=14 animals. (E-F) Activity of single neurons recorded in the superficial (top) and deep superior colliculus (bottom). WT: n=6, ZH3: n=6. Left: (E) Average z-scored neuronal activity during two consecutive looming stimuli. Middle: Cumulative distribution of peak responses of the same neurons. Right: Area under the curve of each response. (F) Contains responses from the same neurons as in E but during a dimming disk stimulus. (G-I) Bayesian model fitted to neuronal data. (G) Average coefficients describing responses to looming and dimming stimuli in the superficial superior colliculus for a model with no genotype information (left) and grouping by genotype (middle). On the right, dispersion of individual mice averages around the total mean in the model with grouping by mouse. (H) Expected log-predictive density difference between the model with grouping by genotype and models with no genotype grouping or grouped by mice. (I) Same as G, for the deep superior colliculus. Statistics were performed using DABEST permutation tests on panels E-F and probability of direction on panels G-I. *p ≤ 0.05. **p ≤ 0.01. ***p ≤ 0.001. ****p ≤ 0.0001, n.s. not significant.

*In vivo* brain recordings revealed neural alterations in the midbrain that correlated with increased fear responses. We presented the same danger-mimicking visual looming stimuli to head-fixed mice while simultaneously recording responses in the superior colliculus, which is the main visuo-motor hub that translates aversive and appetitive stimuli into innate behavioral reactions. In response to a black looming stimulus, we found stronger responses both in neurons in the superficial and deeper layers of the superior colliculus of ZH3 compared to WT mice (Figure 5E). Similarly to the increased behavioral reactions to dimming disks, we also found stronger neural responses in ZH3 mice during this stimulus (Figure 5F). We further analyzed the neural responses to visual stimuli using Bayesian hierarchical models. A model that grouped neurons by genotype confirmed a robust difference between genotypes for looming responses in the deep superior colliculus but not in the superficial superior colliculus (Figure 5G-H). Grouping neurons based on the mouse identity instead did not result in an improvement in the model’s predictive power, showing that the variability in responses across neurons is dominated by the genotype and not by random variability between animals (Figure 5I). In summary, our results suggest that PrP^C^ deficiency affects specific neural computations, enhancing the encoding of both survival-critical and neutral stimuli resulting in stronger behavioral reactions.

## DISCUSSION

### PrP^C^ deficiency destabilizes synaptic network activity

Rather than examining a single experimental level, our data provide a multiscale view of neuronal network alterations associated with chronic PrP^C^ deficiency, encompassing cortical cultures examined at multiple stages of *in vitro* maturation and adult brain circuits. In this study, in fact, we observed that loss of PrP^C^ disrupts the ability of neurons to maintain a stable and fine-tuned network across two independent PrP null mouse models (FVBN-*Prnp*^ZH1/ZH1^ and C57BL/6J-*Prnp*^ZH3/ZH3^ PrP^C^ knockout mice). We used these specific lines since the former is the most widely used line in prion research, while the latter represents the cleanest genetic model currently available: despite differences between the two models, several synaptic and network alterations showed similar patterns across genetic backgrounds, whereas other molecular changes were model specific.

Mature PrP^C^ deficient networks showed less frequent (but a trend towards larger) population events, together with increased pairwise rate correlation, both *in vitro* and *in vivo.* These data are consistent with enhanced network excitability and seizure susceptibility previously reported in PrP null mice ^33,52^. These findings indicate a general effect of PrP^C^ deficiency on neuronal circuits, shifting networks toward a context in which transitions into highly active states become rarer, but when they do occur, activity is abnormally intense and synchronized. Building on previously suggested roles for PrP^C^ in controlling excitability and network dynamics ^31,33^, our findings postulate the view that PrP^C^ acts primarily as a stabilizer of synaptic network activity, rather than simply enabling neurons to fire. Together, these observations are consistent with a contribution of PrP^C^ to the regulation of neuronal network activity, although the mechanisms underlying these alterations remain to be established.

### Molecular correlates of synaptic effects associated with PrP^C^ deficiency

Our molecular analysis indicates that PrP deficiency is associated with a selective remodeling of synaptic components rather than a broad loss of synapses. At later developmental stages, proteomic analyses revealed a broad reduction of presynaptic proteins (including vesicle-release machinery and vesicular transporters) and of postsynaptic components such as PSD95 and glutamatergic receptors in both FVB and ZH3. These findings are in agreement with previous work linking PrP^C^ to synaptic vesicle cycling, Ca²⁺ homeostasis and the maintenance of pre- and postsynaptic structures ^35–37,53,54^. By contrast, changes in inhibitory markers were model dependent, being more evident in mature koFVB cultures than in adult ZH3 cortex, where GAD1-2 were relatively preserved. These molecular changes may be consistent with the altered bursting and increased pairwise rate correlation observed in mature PrP^C^ deficient cultures and in adult cortex. In a network with fewer effective synapses and reduced postsynaptic scaffolding, the recruitment of large population events may be less likely ^55,56^, as reproduced in a minimal mathematical model of an E/I network (Fig. S8). This is consistent with the reduction in PSD95, VGLUT1, SYT1, and CPLX1-2 observed across both PrP^C^ deficient models, suggesting reduced glutamatergic drive, altered postsynaptic organization, and altered presynaptic vesicle release. Importantly, the molecular alterations measured here should not be interpreted as a direct explanation of the earlier network phenotype and may instead represent later emerging consequences or adaptations associated with chronic PrP^C^ deficiency. Moreover, as both PrP deficient lines used in this study lack PrP throughout development, we cannot exclude that some of the molecular alterations observed in knockout neurons and adult tissue reflect developmental compensation rather than direct, acute consequences of PrP loss. This possibility is particularly relevant when interpreting synaptic protein changes, as chronic absence of PrP may trigger adaptive remodeling of excitatory and inhibitory synaptic components during network formation and maturation.

### Shared and model-specific consequences of PrP^C^ loss

At the functional level, the two PrP-deficient models showed partially overlapping alterations in network activity. Mature koFVB cultures displayed less frequent network-wide bursts together with increased pairwise rate correlations. ZH3-derived cortical cultures showed a comparable pattern in burst occurrence under the same recording conditions, although these differences did not reach statistical significance in the small cohort. Consistently, adult ZH3 recordings revealed related alterations in network activity in vivo, including a region-dependent reduction in burst occurrence and increased correlation between cortical neurons. Thus, despite differences in genetic background and experimental context, PrP^C^ deficient networks appear to be characterized by less frequent population events and stronger coordination of neuronal activity.

A similar convergence between the two models was observed at the protein level, mostly suggesting that impaired maintenance of excitatory synapses is a robust consequence of PrP^C^ loss, particularly at late developmental stages, when circuits must remain stable while still retaining the capacity to adapt ^34,57^. However, the two models differed transcriptionally. In line with the original description of the co-isogenic ZH3 line ^32^, RNA-seq from ZH3 cortical cultures at DIV23 showed only modest transcriptomic changes, largely restricted to *Prnp*, whereas the FVB model displayed broader (even if still minimal) pathway-level dysregulation. This mismatch, limited changes at the RNA level in ZH3 but reproducible synaptic protein deficits in both models, raises the possibility that PrP^C^ loss impacts synapses mainly through post-transcriptional mechanisms, altered protein stability/turnover, or local synaptic regulation that is not well captured by bulk RNA-seq ^58^. Genetic background may also influence how strongly compensatory programs are engaged: koFVB networks may mount broader transcriptional responses, while ZH3, being co-isogenic, may show a more constrained transcriptomic profile even when the protein-level endpoint converges.

Furthermore, in koFVB cultures, reductions extended across all proteins investigated at maturity, whereas ZH3 tissue showed a more selective convergence on core excitatory synaptic characters such as VGLUT1 and PSD95, and on the presynaptic machinery with SYT1 and CPLX1-2, while other related components remained intact. This likely reflects a combination of factors, including developmental stage and experimental context (maturing networks *in vitro* versus stable adult circuits *in vivo*), differences in cellular composition and circuit architecture between dissociated cultures and intact cortex, and the action of homeostatic mechanisms *in vivo* that may partially buffer presynaptic deficits while leaving excitatory synapse scaffolding and transporter expression particularly vulnerable ^55,59^. Together, these similarities and differences support a model in which PrP^C^ helps sustain excitatory synapse integrity across contexts, while the extent of downstream remodeling depends on genetic background and the capacity for compensation.

### PrP^C^ deficiency effects on states and traits in adulthood

In vivo brain recordings and behavior assays confirm that network alterations due to PrP^C^ deficiency are traceable into adulthood and affect specific neural computations. Previous reports on neural and behavioral changes have shown that a lack of PrP^C^ impairs memory, attention, operant learning and long-term potentiation ^18,33,60,61^. Here, we have expanded these findings to other types of learning such as adaptation to stimulus repetitions and environmental context where we found a similar impairment in KO mice. In addition, effects on anxiety in elevated plus maze and open field tests were reported in the ZH1 mouse line ^62^. Previous studies also found an effect on novel object recognition ^33^. However, we argue that these experiments were performed under high stress conditions and that this assay, when performed under low stress, is better considered as a test for novelty curiosity and exploratory behavior than for learning. In our hands, we found no significant differences between WT and ZH3 mice in any of our assays testing for changes in behavioral state, including exploration/curiosity (novel object recognition task), anxiety (open-field, plus maze, dark-light chamber, marble burying) or sociability (social interaction task).

### PrP^C^ deficiency leads to behavioral and neuronal overreaction to danger

While we found no effect on state, PrP^C^ deficient mice displayed more and earlier innate evoked reactions to overhead danger. Previous work has found that ZH1 mice show decreased freezing in response to a real snake ^61^ but increased fear learning ^62^. In ZH3 mice, we found increased escape behavior not only to looming threat, but also to a dimming disk stimulus which is considered an ethologically irrelevant stimulus, normally producing few reactions and weak neuronal activity ^49,51,63^. Importantly, these behavior changes correlated with stronger neuronal responses to the same stimuli in the superior colliculus, the main visuo-motor hub implicated in innate aversive behaviors. A significant difference in the activation of PrP^C^ deficient superior colliculus neurons in its superficial layers (implicated in saliency detection) and even more so in its deeper layer (implicated in sensory integration and escape mediation) is supported by our computational model. In fact, genotype explained the differences in neuronal firing better than mouse identity. This neuronal and behavioral overreaction is consistent with circuits that spend much of their time in a relatively low-activity state but, once engaged, respond with overly synchronized and prolonged high-amplitude events.

### PrP^C^ as a tuned regulator of synaptic and network balance

In combination, our molecular, *in vitro,* and *in vivo* experiments provide evidence that PrP^C^ contributes to the maturation and stabilization of neuronal networks, affecting both their electrophysiological behavior and molecular architecture. During early stages, PrP^C^ promotes neurite growth and the establishment of balanced synaptic connections, facilitating the emergence of coherent activity patterns ^5,64,65^. However, in its absence, no clear network phenotype was evident at the earlier stages examined. At maturity, instead, alterations in synaptic maintenance, translational control, and excitatory spine stability may contribute to reduced excitatory synapse density and altered network dynamics. Regardless of the precise mechanisms, our data suggest that the absence of PrP^C^ affects the fine-tuning of network development, leading to delayed maturation, altered bursting, and changes in pairwise rate correlation. Moreover, *in vivo* brain recordings and behavioral assays show that related networks alterations are present into adulthood and across different genomic backgrounds. The observation of similar patterns in both FVB and ZH3 models, and *in vitro* and *in vivo*, argues that impaired control of network dynamics is a characteristic consequence of PrP^C^ loss rather than a peculiarity of a single preparation or experimental setup, consistent with the phenotype observed in prion and other neurodegenerative diseases.

### Implications for prion disease treatment strategies

These results could have direct implications for strategies that aim to lower PrP^C^ to treat or prevent prion diseases. Genetic deletion of PrP^C^ protects against prion replication and confers resistance to scrapie ^48^, and several pharmacological and genetic approaches to reduce PrP^C^ expression are under active development as disease-modifying therapies ^44–46^. While targeting PrP^C^ remains a promising approach to counteract prion diseases, our data reveal that chronic loss of PrP^C^ could cause significant neuronal network impairments, especially at mature stages and in adulthood. Although such approaches may be less critical in sporadic prion diseases, where the late onset of pathology may have limited impact on the physiological role of the protein, they could become particularly relevant for prophylactic interventions in genetic prion diseases during adulthood, as well as in iatrogenic forms of the disease. Consequently, researchers working on therapeutic strategies should consider the risk of destabilizing normal synaptic and network functions, while balancing the benefits of PrP^C^ removal, particularly in the context of aging or neurodegeneration. It remains to be tested whether ablation of prion protein in the mature brain, after having fully developed in the presence of the protein, replicates the effects we describe here. In fact, the models used in the present study undergo complete, lifelong germline deletion of PrP, whereas therapeutic approaches are expected to reduce PrP after development and may result in only partial lowering. Future studies tackling this question are enabled by recent developments in prion protein targeting viral vectors ^66,67^.

In summary, by integrating longitudinal MEA recordings, synaptic protein profiling, adult *in vivo* electrophysiology, and behavioral assays across two PrP null models and multiple developmental stages, we provide a multilevel characterization of the physiological consequences of chronic PrP^C^ deficiency. This integrated approach identifies convergent alterations in synaptic organization and network dynamics that are accompanied by altered stimulus evoked behavioral responses, and highlights an important physiological constraint for the design of therapies targeting PrP^C^.

## RESOURCE AVAILABILITY

### Lead contact

Further information and requests for resources and reagents should be directed to and will be fulfilled by the Lead Contact.

### Materials availability

This study did not generate new unique reagents.

### Data and code availability

RNA-seq, *in vitro* electrophysiology, *in vivo* electrophysiology and behavior preprocessed datasets are deposited in a public repository accessible via https://doi.org/20.500.12928/FMSJL3. Additional analysis scripts will be shared on reasonable request.

## AKNOWLEDGMENTS

The authors thank Sequentia Biotech (Barcelona, Spain) for generating the RNA-sequencing data presented in this study, the technical staff and the Animal House staff in SISSA for their support throughout the project. We also would like to thank Maria del Carmen Costas Ferreira for providing the graphical abstract. Authors acknowledge generous funding from the European Research Council (ERC) (Grant agreement No. 101075848 to K.R.), the European Innovation Council (GA. 101070908, “CROSSBRAIN”, to M.G.), the National Science Foundation (award 2515404, to M.G.), the National Recovery and Resilience Plan (NRRP), Mission 4, ’Education and Research’ Component 2, ’From research to Business’ Investment 3.1 – Call for tender 3264 of 28/12/21 of MUR - Italian Ministry of Universities and Research, funded by the EU – NextGenerationEU (project IR0000011, Concession Decree 117 of 21/6/22 adopted by MUR, CUP B51E22000150006, “EBRAINS-Italy” to M.G.). The authors also wish to thank the generous fundings from Target ALS Foundation (BM-2022-C3-L2) and the PNRR Mission 4, Component 2, Investment 1.3_PE_00000015_AGE-IT - spoke 2 - fincanced by the European Union – NextGenerationEU – CUP: G93C22001070006 (both awarded to G.L.). The views and opinions expressed are solely those of the authors and do not necessarily reflect those of the European Union, nor can the European Union be held responsible for them.

## AUTHOR CONTRIBUTIONS

A.B (Conceptualization, Data curation, Formal analysis, Investigation, Methodology, Validation, Visualization, Writing – original draft, Writing – review & editing), A.D.C (Conceptualization, Data curation, Formal analysis, Investigation, Methodology, Validation, Visualization, Software, Writing – original draft, Writing – review & editing), C.L (Conceptualization, Data curation, Formal analysis, Investigation, Methodology, Validation, Visualization, Writing – original draft), V.P (Formal analysis, Investigation, Methodology, Validation, Visualization), G.P (Formal analysis, Investigation, Methodology, Validation, Visualization), B.E.P.M (Formal analysis, Investigation, Methodology, Writing – original draft), B.P (Investigation, Methodology), M.Z. (Conceptualization, Visualization, Supervision, Writing – review & editing), L.C. (Conceptualization, Visualization Supervision, Methodology), L.Z (Investigation, Methodology, Formal analysis, Validation, Visualization), L.S. (Conceptualization, Investigation, Methodology), E.P (Conceptualization, Supervision, Project administration, Writing – review & editing), M.G. (Conceptualization, Supervision, Project administration, Funding acquisition, Writing – original draft , Writing – review & editing), K.R. (Conceptualization, Supervision, Project administration, Data curation, Formal analysis, Funding acquisition, Visualization, Writing – original draft, Writing – review & editing), G.L. (Conceptualization, Supervision, Project administration, Funding acquisition, Writing – original draft, Writing – review & editing).

## STAR METHODS

Key Resources in **Table 1** below

**Table 1.**
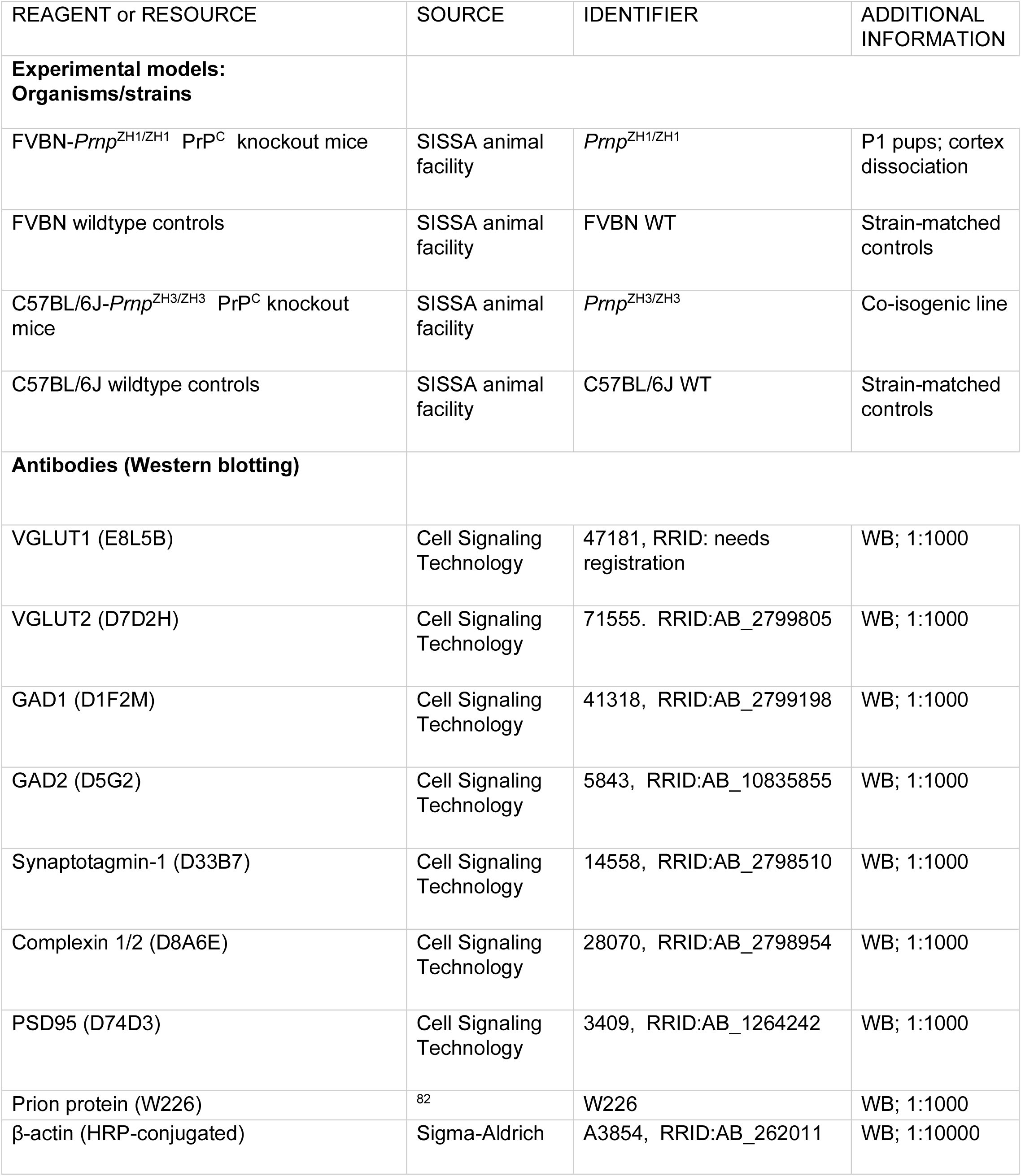
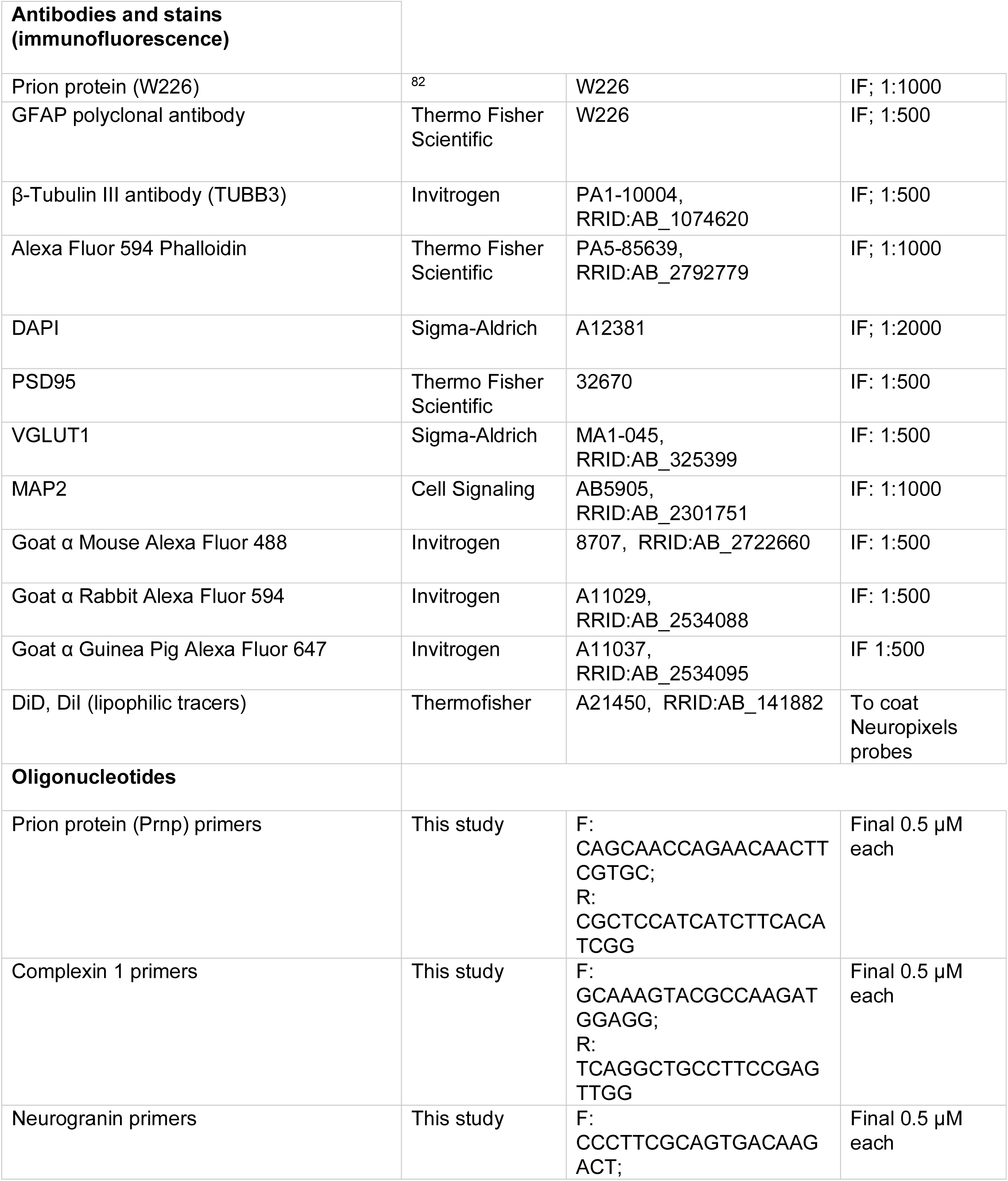
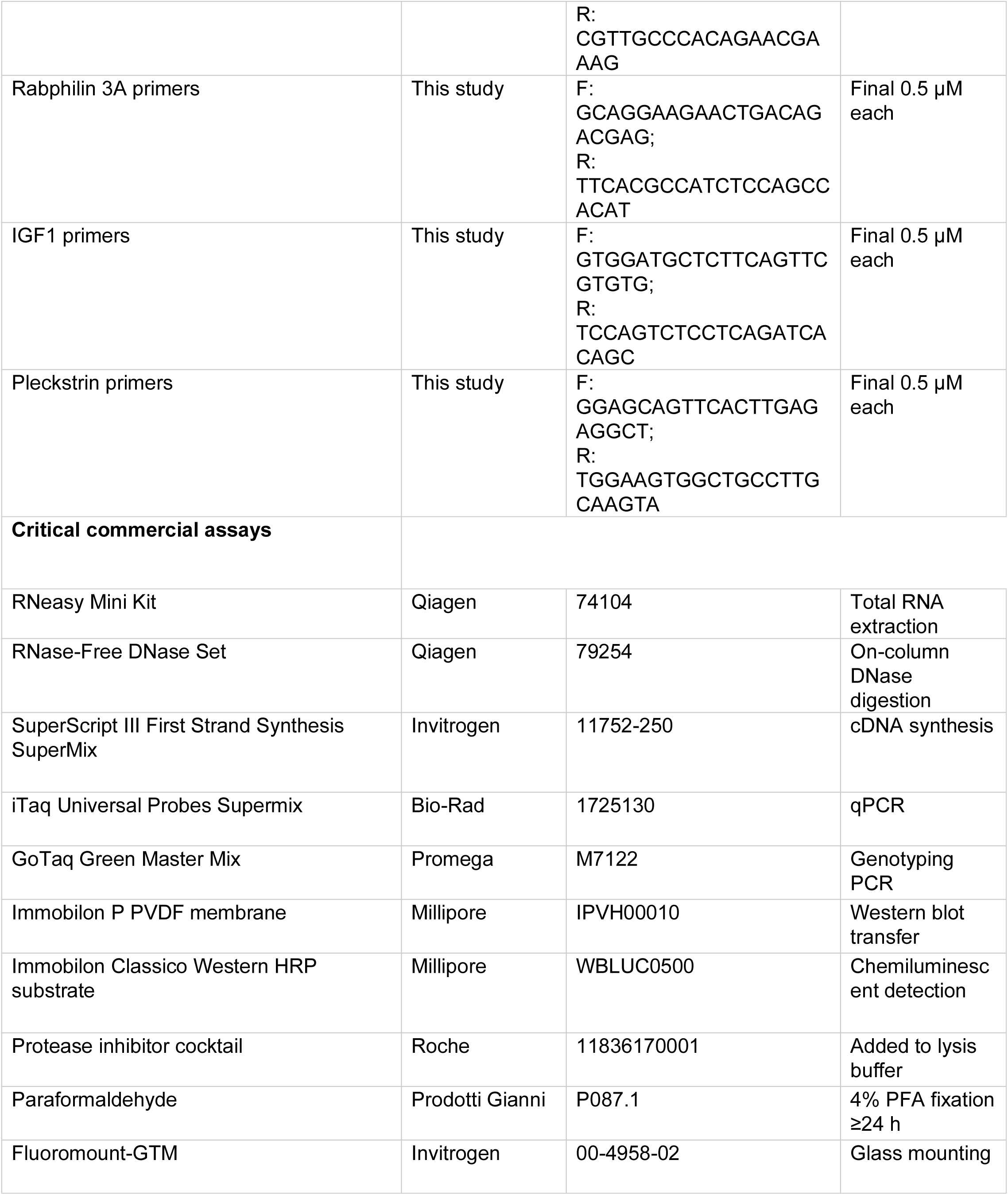
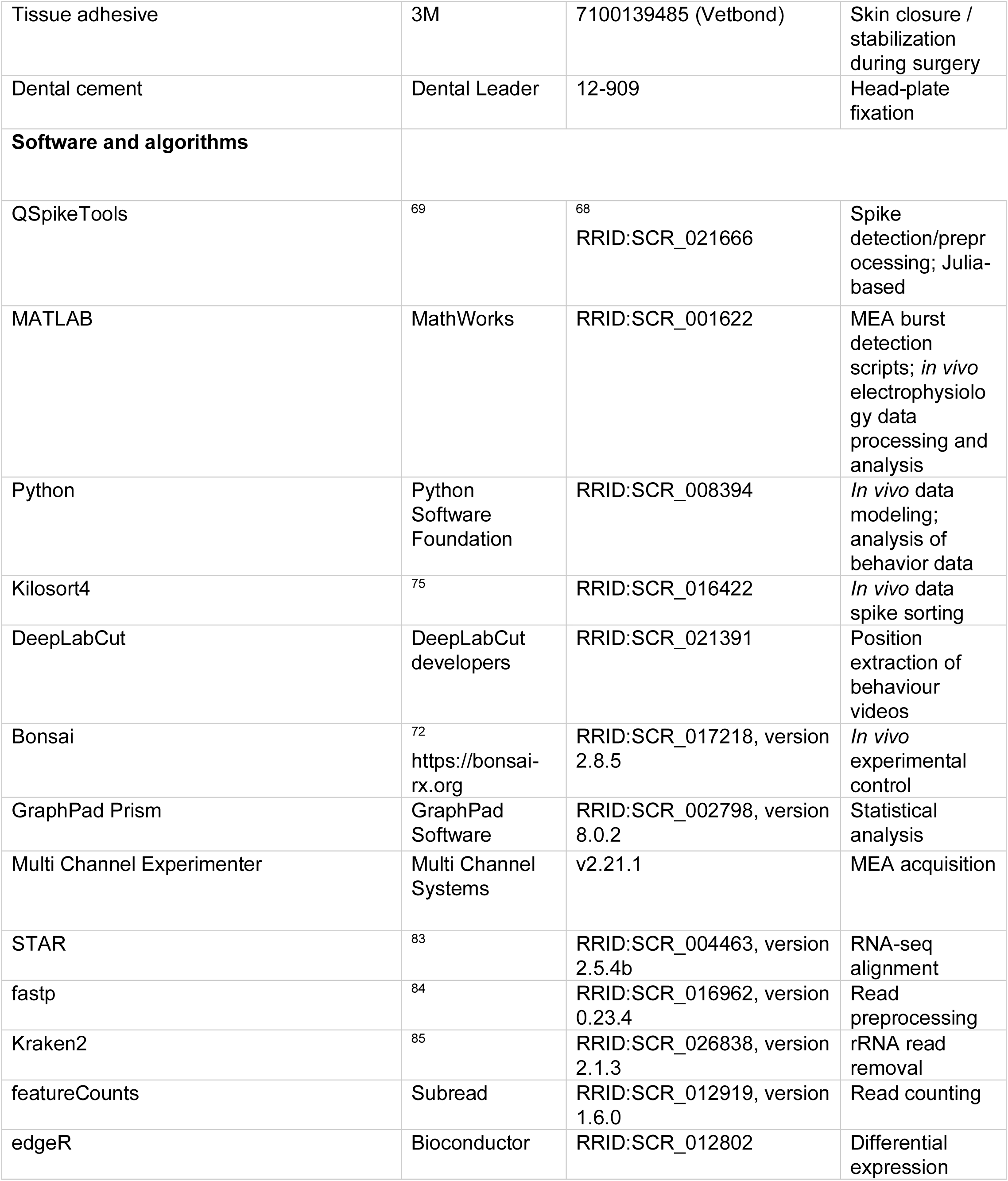
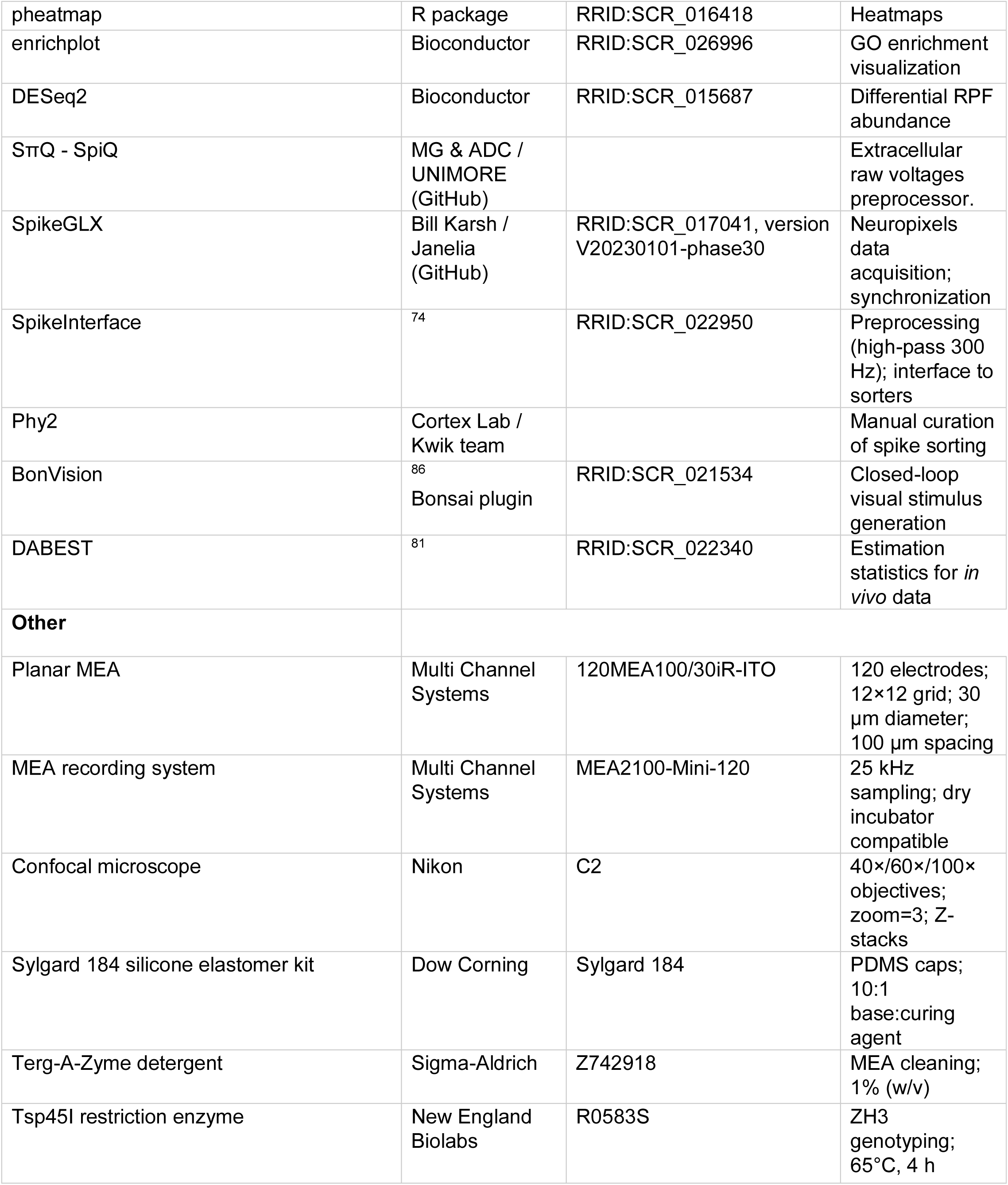
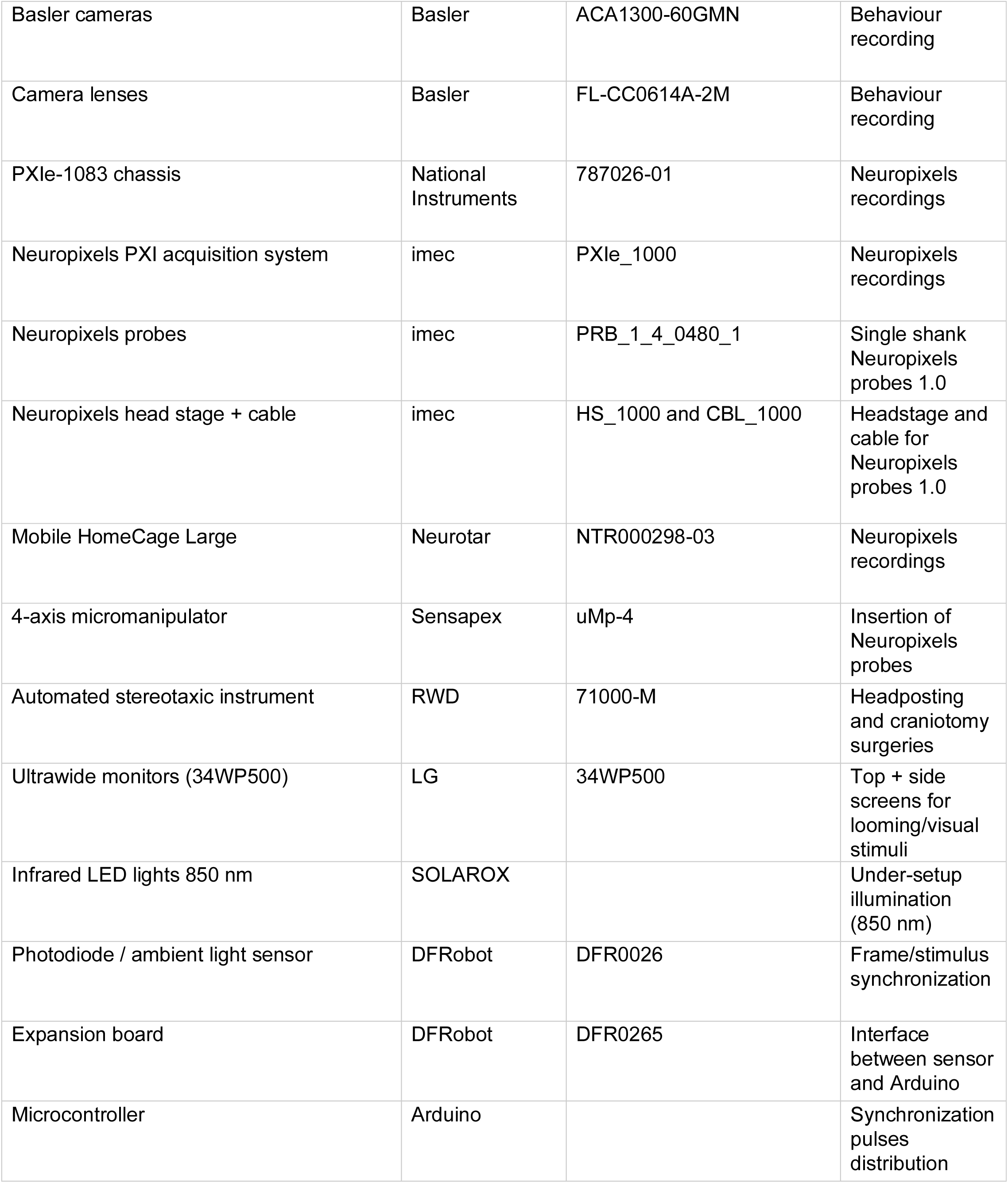
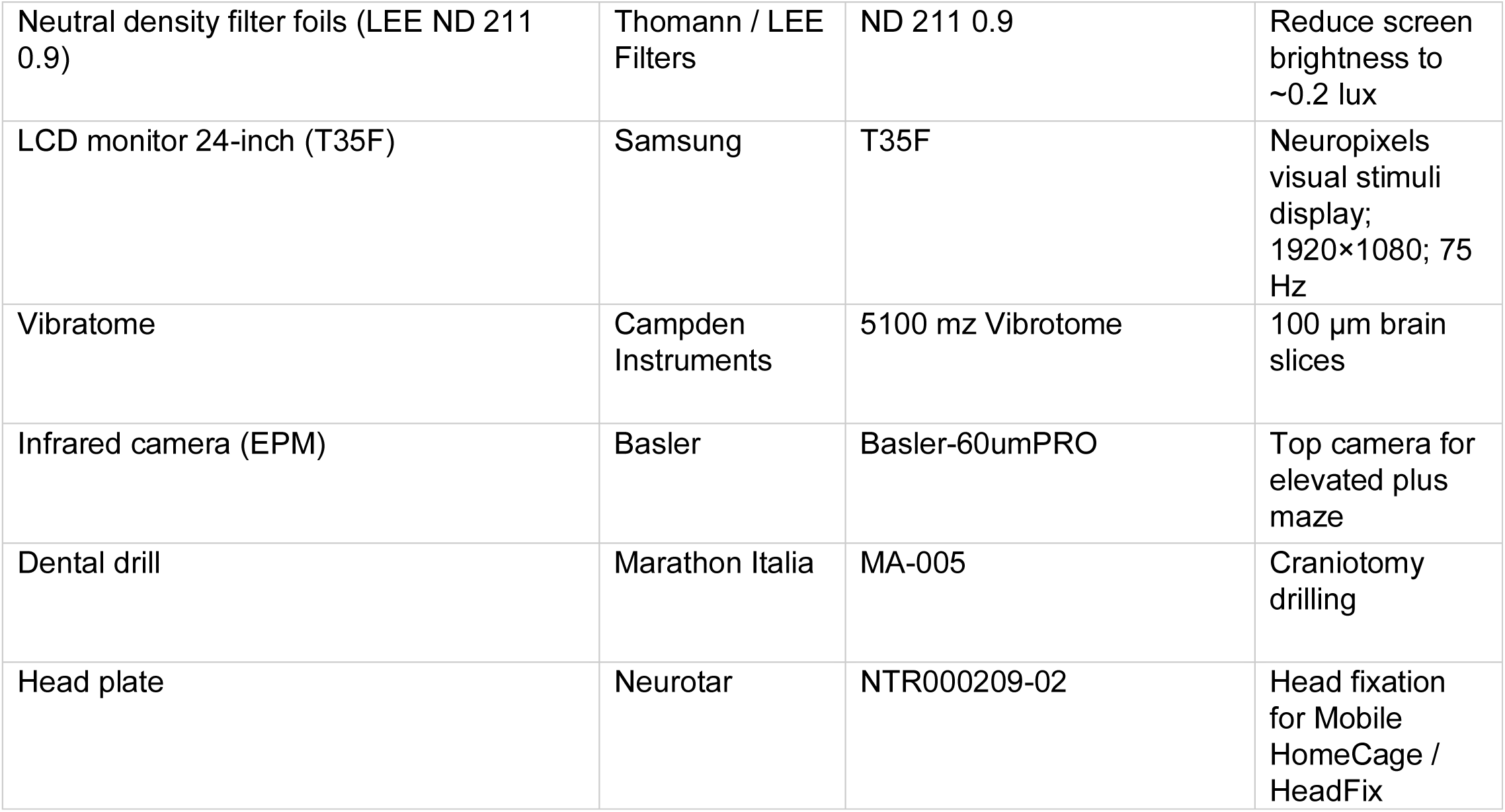
Summary of all resources used in this study.

### Mice

All animal procedures were approved by the Italian Ministry of Health and local veterinary authorities (OPBA, SISSA) and performed in accordance with Italian decree 26/2014 and EU directive 2010/63/EU, with efforts to minimize animal number and suffering. Experiments with adult C57BL/6J-Prnp^ZH3/ZH3^ (referred to as ZH3) were approved by the Italian Ministry of Health with authorization number 703/2023-PR.

Primary cultures were prepared from postnatal day 1 (P1) pups from FVBN-PrnpZH1/ZH1 prion protein (PrP) KO mice and strain-matched wildtype controls. Also, the co-isogenic C57BL/6J-Prnp^ZH3/ZH3^ line was used in this study.

### Solutions and media

Solutions were prepared as follows:

### Primary cortical neuron culture

Well plates were coated with poly-L-ornithine (1:1000 in ddH2O) and MEAs with 1 mL 0.1% polyethyleneimine (PEI) 24 h before dissection (overnight at 37°C).

P1 pups were sacrificed by decapitation. Brains were dissected in dissection medium (See Table 2 for media compositions). Cortices were chopped and enzymatically dissociated in digestion medium containing trypsin (6000 U/mL) and DNase (1560 U/mL) for 5 min at 37°C (5% CO2). Digestion was quenched by washing with dissection medium and incubating with trypsin inhibitor (1000 U/mL, in dissection medium) for 10 min at 4°C. Tissue was washed in dissection medium containing DNase (1248 U/mL) and mechanically dissociated. Cells were centrifuged at 100×g for 5 min, resuspended in culture medium, and viable cells were counted using a Bürker counting chamber.

**Table 2.** Composition of media necessary for neuronal culture dissociation.

| Solution | Composition |
| --- | --- |
| Culture medium | 92% Neurobasal-A, 5% horse serum, 1% GlutaMAX, 2% B27, gentamycin (50 mg/mL). |
| Dissection medium | Hanks' solution (Ca<sup>2+</sup>/Mg<sup>2+</sup>-free; 9.52 g/L) + 4.2 mM NaHCO<sub>3</sub>, 12 mM HEPES, 33 mM D-glucose, 200 μM kynurenic acid, 25 μM APV, 5 μg/mL gentamycin, 300 mg/L BSA, 12 mM MgSO<sub>4</sub>; pH 7.3; filtered; stored at 4°C. |
| Digestion medium | 137 mM NaCl, 5 mM KCl, 7 mM Na<sub>2</sub>HPO<sub>4</sub>, 25 mM HEPES, 4.2 mM NaHCO<sub>3</sub>, 100 μM kynurenic acid, 25 μM APV; pH 7.4. |

For MEA experiments, 1.8 × 10^6 cells in 1 mL were plated per MEA (6500 cells/mm^2) and sealed with sterile PDMS caps. For biochemical assays, 1.5 × 10^6 cells in 2 mL were plated per coated 6-well. For immunofluorescence, 1 × 10^5 cells in 1 mL were plated on 15 mm coverslips. Cultures were maintained at 37°C and 5% CO2; 50% medium was replaced 72 h after plating and every 48 h thereafter.

### Genotyping

Tail biopsies were digested overnight at 55°C in tail lysis buffer containing proteinase K (25 µL/mL). Samples were vortexed, centrifuged (400×g, 10 min, room temperature), and supernatants were diluted 1:15 in sterile water.

FVBN-PrnpZH1/ZH1 genotyping PCR used GoTaq Green Master Mix (per sample: 7.5 µL mastermix, 1.1 µL ddH2O, 1 µL each primer RK1/RK2/RK3 at 10 µM, 0.4 µL DMSO). Cycling: 94°C 5 min; 35 cycles of 95°C 40 s, 58°C 50 s, 72°C 1 min 30 s; final extension 72°C 7 min; hold 4°C. Products were resolved on 1.2% agarose in 0.5× TBE with Midori Green; electrophoresis 90 V 15 min then 120 V 30 min.

C57BL/6J-Prnp^ZH3/ZH3^ PCR used ZH3-F/ZH3-R (20 µM each), with cycling: 94°C 2 min; 35 cycles of 94°C 30 s, 58°C 30 s, 72°C 45 s; final extension 72°C 5 min. PCR products were digested with Tsp45I (NEB) at 65°C for 4 h (CutSmart buffer) and resolved on 2.3% agarose in 0.5× TBE with Midori Green.

### Microelectrode array recordings

Planar MEAs (120MEA100/30iR-ITO; 120 ITO electrodes in a 12×12 grid; 30 µm electrode diameter; 100 µm spacing) were used. MEAs were sterilized in 70% ethanol (30 min), rinsed three times in sterile Milli-Q water, dried, and coated overnight at 37°C with freshly prepared 0.1% PEI (1 mL/MEA). MEAs were rinsed three times and dried before seeding. After use, MEAs were cleaned overnight at 4°C in 1% (w/v) Terg-A-Zyme, washed, dried, then filled with sterile Milli-Q water and stored at 4°C in the dark; water was exchanged at least monthly. Caps were fabricated from Sylgard 184 mixed 10:1 (base: curing agent), rested 30 min at room temperature, poured into molds, and cured 1 h at 100°C. Caps were ethanol-sterilized and dried; caps were reused 1–2 additional times depending on condition.

Recordings were acquired with an MEA2100-Mini-120 system (Multi Channel Systems) in a dry incubator environment (37°C, 5% CO2). Capped MEAs were mounted using a sandwich contact arrangement. Signals were acquired with Multi Channel Experimenter at 25 kHz.

Spontaneous activity was recorded for 30 min at DIV10, DIV17, and DIV23 (52 MEAs across culture sessions). Cultures were stabilized for 10 min prior to acquisition to reduce inter-variability. Recordings were included in the analysis if at least 75% of electrodes (≥90/120) exhibited activity.

### MEA signal processing and network metrics

Raw recordings (.msrd) were analyzed with QSpikeTools ^68^ . Files (>20 GB) were converted to .h5 and processed on the SISSA Ulysses cluster with channel-wise parallelization.

Spike detection was performed per channel using an unsupervised detector with acausal filtering ^69^, yielding per-electrode spike-time lists. Active electrodes were defined as electrodes with firing rate > 0.02 Hz. Population spiking rate was computed as total spikes divided by (active electrodes × recording time). Burst detection was performed from spike-time histograms (5 ms bins) using a threshold-crossing method with a threshold set to 15% of the number of active electrodes. Burst onset/offset were defined by the first 10 consecutive bins with <2 spikes before/after the burst peak. Cross-correlation was computed for each electrode pair by counting spikes from train Y within ±T of spikes in train X using T = 300 ms and Δt = 3 ms. Cross-correlograms were normalized by the square root of the product of spike counts. Significance was assessed against 100 surrogate correlograms generated by shuffling inter-spike intervals.

### MEA data statistical analysis

For total spike counts, which yielded one value per MEA, normality was assessed using the Shapiro– Wilk test. Normally distributed data were compared between genotypes using two-tailed unpaired *t*-tests with Welch’s correction, whereas non-normally distributed data were analyzed using two-tailed Mann–Whitney U tests. For burst-related parameters (inter-burst interval, burst amplitude, burst onset slope, and burst duration) and pairwise cross-correlation indices, repeated observations were nested within individual MEAs and were therefore analyzed using a clustered Wilcoxon rank-sum test implemented with within-cluster resampling permutation (WCRP) as described by Follmann and Fay ^70^. The MEA was treated as the independent cluster, and genotype labels were permuted only at the whole-MEA level, thereby preserving the dependence among observations within each recording. Genotype permutations were exhaustive when computationally feasible and otherwise based on 100,000 Monte Carlo permutations. *P* values were adjusted for the three developmental-stage comparisons (DIV10, DIV17, and DIV23) within each outcome using the Holm procedure. Effect sizes for clustered analyses were expressed as cluster-weighted rank-biserial correlations (*r_rb_*).

### Western blotting

Primary neuronal cultures were harvested at DIV10, DIV17, and DIV23. Cells were rinsed twice in PBS (pH 7.4) and lysed in 50 mM Tris (pH 7.4), 250 mM NaCl, 5 mM EDTA, 50 mM NaF, 1 mM Na3VO4, 1% NP-40, 0.02% NaN3, supplemented with protease inhibitor cocktail. Lysates were centrifuged at 5900×g (4°C) and supernatants stored at −20°C. Protein concentration was determined by BCA assay. Samples (20 µg) were prepared in 4× loading buffer (10% glycerol, 50 mM Tris-HCl, 2% SDS, bromophenol blue, freshly added 200 mM DTT) and boiled for 10 min at 100°C. Proteins were resolved on mini-PROTEAN TGX gels and transferred to PVDF membranes for 2 h at 300 mA (4°C). Membranes were blocked in 5% non-fat milk in TBS-T and incubated overnight at 4°C with primary antibodies (see Key Resources Table). After washes, membranes were incubated with HRP-conjugated secondary antibodies for 1 h at room temperature and developed with chemiluminescent substrate. Images were acquired on an iBright imaging system (ThermoFisher). β-actin-HRP was used for normalization.

### Immunofluorescence

Dissociated cortical cells were plated on 15 mm coverslips at 500 cells/mm^2. Coverslips were coated overnight at room temperature with poly-L-ornithine (150 µL), rinsed 3–4 times with sterile water, and cells were allowed to attach for 30 min before adding 2 mL culture medium. For fixation, cells were washed three times in 1xPBS (5 min each) and fixed in 4% PFA in 1xPBS (pH 7.2) for 30 min at room temperature, and then washed again 3 times in 1xPBS. For permeabilization, cells were incubated in Blocking Solution with 10% Normal Goat Serum and 0,1% Triton for one hour. Later, cells were incubated with primary antibodies (see Table 1) diluted in 1:5 of Blocking Solution and stored at 4°C overnight.

The following day, cells were washed with 1× PBS three times for 5 minutes each. Then they were incubated with secondary antibodies (see Table 1) in 1:5 of Blocking Solution in 1× PBS for one hour and then washed three times for 5 minutes each with 1×PBS. The coverslips were then mounted on a glass-slides sealed with Fluoromount-GTM, letting them dry at room temperature in a dark box.

### Image acquisition and processing

Images from immunofluorescence experiments on cell cultures were acquired using a Nikon A1X confocal microscope with different objectives at the SISSA microscopy facility. 40× images were used for neuronal quantification and 60× oil immersion images were used for synaptic quantification. Each image was acquired using four lasers with different wavelengths: FITC (488 nm), TRITC (564 nm), Cy5 (647 nm), and DAPI (408 nm). For the 60× objective, z-stacks consisting of 75 slices with a step size of 0.2 µm were acquired. Images were analysed using Volocity software.

### Synaptic quantification

Nuclei were identified in the DAPI channel using the standard deviation method and filtered by excluding objects with a volume smaller than 20 µm³. Neurons were identified in the TRITC channel using the standard deviation method, and objects with a volume smaller than 50 µm³ were removed.

Presynaptic elements were identified in the FITC channel based on intensity. To refine the measurement, objects were filtered by removing those with a volume below 0.1 µm³, and touching elements were separated using multiple object size guides, ranging from 10 µm³ to 3 µm³.

Postsynaptic elements were identified in the Cy5 channel using the intensity method and filtered by excluding objects larger than 10 µm³ and smaller than 0.01 µm³. Touching objects were separated using the “separate touching objects” function with multiple object size guides, ranging from 100 µm³ to 5 µm³.

Presynaptic and postsynaptic elements within neurons were quantified by counting vGlut1 and PSD95 objects, respectively, that interact with MAP2 objects.

For active synapses, only objects in the FITC channel that interact with those in the Cy5 channel were considered as total synapses, while active synapses within neurons were defined as PSD95 and vGlut1 objects that are in contact with each other and also with MAP2 objects.

### Neuronal quantification

Nuclei were identified in the DAPI channel using the standard deviation method and filtered by excluding objects with a volume smaller than 20 µm³. Neurons were identified in the TRITC channel using the standard deviation method, and objects with a volume smaller than 50 µm³ were removed. Neurons were then counted considering only MAP2 objects that were in contact with nuclei. The number of neurons in each field was normalized to the surface area of the field, obtaining neuronal density for WT vs KO conditions.

RNA extraction, reverse transcription, and RT-qPCR Total RNA was extracted from primary cortical cells using RNeasy Mini Kit with on-column DNase digestion (RNase-Free DNase Set). RNA was quantified by Nanodrop (DeNovix) and stored at −80°C. cDNA was prepared using SuperScript III First Strand Synthesis SuperMix. qPCR was performed with iTaq Universal Probes Supermix on a CFX96 Touch system using 10 ng/µL template. Primer sequences are reported in the Key Resources Table (final 0.5 µM each). Cycling: 95°C 30 s; 40 cycles of 95°C 15 s and 60°C 30 s; hold 4°C. No-RT controls and water controls were included. Expression values were normalized to β-actin and calculated using the 2^(-ΔΔCt) method^71^.

### RNA sequencing and differential expression analysis

RNA integrity (RIN) was assessed on an Agilent BioAnalyzer 2100. Libraries were preprocessed using fastp (v0.23.4). rRNA reads were removed using Kraken2 (v2.1.3). Reads were aligned to Mus_musculus.GRCm39.105 using STAR (STAR-2.5.4b) with end-to-end alignment (alignEndsType EndToEnd). Gene-level quantification used featureCounts (v1.6.0) counting reads with minimum mapping quality 30. Features were filtered (≥10 counts and present in at least two samples). PCA and heatmaps (pheatmap) were used for exploratory analysis. Differential expression analysis was performed with edgeR (FDR < 0.05). GO enrichment analysis for up- and down-regulated genes was performed using enrichplot, visualized with dotplots/barplots/enrichment maps across GO ontologies.

## IN VIVO BEHAVIOUR EXPERIMENTS

### Mice

*In vivo* experiments were performed with C57BL/6J-*Prnp*^ZH3/ZH3^ homozygote knockout and wildtype littermates. Animals were housed in the SISSA facility until 1-2 weeks before experiments. Then, they were kept in housing cabinets in the experimental rooms to acclimate to the environment. Mice were kept under 12h/12h dark/light conditions with food and water *ad libitum* and humidity of 45-65%. At the beginning of behavioral and electrophysiological experiments, mice were 12 weeks old. Animals of both sexes were used for experiments.

### Experimental set-up for visual stimuli, open-field assay, novel object and social interaction task

Behavioral experiments to determine responses to a visual threat stimulus were carried out in a customized rectangular arena (83 cm W x 35 cm D x 37 cm H) with a separated entrance area attached to one short side (35 cm W x 35 cm D x 37 cm H). The base of the arena was made of IR transmissive red plexiglass while the top and one of the long sides consisted of two monitors (34WP500 Ultrawide Monitor LG) where visual stimuli were programmed and presented with Bonsai Software^72^. Deep red lights (SOLAROX LED 850 nm) were placed under the setup. In the left corner of the arena was placed a triangular-shaped shelter made of red Plexiglas (20 cm W x 10 cm H). Behavioral responses were recorded by three infrared cameras (Basler ace acA1300-60gmNIR; lens FL-CC0614A-2M) at 60 fps placed below the floor and on the top of both the short-side of the arena, respectively. A photodiode (DFRobot Ambient Light Sensor DFR0026) was attached to one of the corners of the top LCD monitor and connected to an Arduino Uno via an expansion board (DFRobot DFR0265), which was used to synchronize individual frames with the stimulus.

### Visual stimuli for behavior assays

Visual stimuli were created using a Bonsai Software plugin, BonVision (a closed-loop visual environment generator, Lopes et al., 2021). Stimuli consisted of (1) a black looming disk on a grey background and expanding from 2 to 40° in 500 ms, followed by 500 ms of grey screen, repeated twice; (2) a dimming disk of a size of 40° which dimmed from grey to black within 500 ms, followed by 500 ms of grey screen, repeated twice. The stimuli were shown on the top screen on the opposite side of the shelter and repeated three times for each session. The screen brightness (grey) was 100 lux to which three neutral density filter foils (Thomann, LEE ND filter 211 0.9 ND) were taped to decrease the brightness to ∼0.2 lux.

### Looming assay

At least three days before the first experimental session each mouse was placed in the arena for 30 min to habituate to the set up. On the testing day mice were moved from the cage into the entrance area of the setup using a transport box which they entered and left on their own. Once the mouse entered the main setup area, experiments were started. After 15 minutes of recording the spontaneous behavior in the arena, the first visual stimulus was triggered as soon as the animal entered the dedicated threat zone on the opposite side of the shelter. The order of stimuli and time between each stimulus (between 60 s and 180 s) were pseudorandomly chosen. After this time, the next stimulus was triggered at the next entry into the threat zone. Each animal completed 2 experimental sessions with at least 7 days between sessions. All the experiments were performed between 9 am and 12 am at Zeitgeber 2-5. After data processing with DeepLabCut (see below), the following parameters were extracted: maximum speed after stimulus, the latency to escape and the percentage of type of response after the stimulus. The maximum speed was calculated as the maximum speed reached by the animal within the first 5 seconds after stimulus onset. The latency instead was measured as the time of escape onset, i.e. the duration between stimulus onset and the moment at which the animal reached a speed of >30 cm/s for at least 0.16 s. The escape and freezing threshold of respectively 30 cm/s and 0.4 cm/s were calculated as the 97.5 and 2.5 percentile of the cumulative distribution of the speed during the 15 min of habituation. Finally, for each animal, behavioral responses were classified into three mutually exclusive categories: "Escaping", "Freezing", and "Not responding". “Escaping” and “Freezing” were defined by a minimum of 20 consecutive frames with speed above the escaping threshold or below the freezing threshold, respectively. If both criteria were fulfilled, the trial was classified as “Escaping”.

### Data processing

Mouse tracking was performed using DeepLabCut (v3.0, Mathis et al., 2018; Nath et al., 2019). The network (ResNet-50 backbone) was trained for 100000 iterations on manually annotated frames to extract the positions of multiple body landmarks, including the center of the body, paws, head, nose, and tail. A consensus across tracked body parts was used to estimate the animal’s center of mass. Custom-written scripts in Python were used to visualize and manually curate tracking data. For each frame, the position of the animal was calculated from the x and y coordinates of the center as √(x² + y²). Instantaneous speed was computed as the frame-to-frame difference in position. Speed values were smoothed using a moving median window of five data points and converted to cm/s based on pixel-to-centimeter calibration derived from the known dimensions of the experimental setup.

### Open field Test

The 15 min of habituation before the looming assay represent a standard Open Field Test. Data was processed using DeepLabCut as described above and the following parameters were extracted: number of entries in the center of the arena, the total distance travelled, the average speed during the session and the time spent under the shelter. The center of the arena was defined as a rectangular area in the center of the setup that measured 18 cm x 47 cm and an entry in the area was counted every time the body center coordinates of the animal crossed the edges of this rectangle. The time spent under the shelter was defined as the total duration during which the animal’s body center coordinates were located within the shelter area. The shelter area was defined as a circular region centered on the shelter, with a radius of 9 cm.

### Dark-Light Test

The test was conducted in the previously described arena divided into two compartments by a black plexiglass partition, with the dark compartment occupying one-third of the arena. The dark compartment was enclosed with black fabric on the lateral walls and upper monitor to minimize light exposure, while the remaining area was brightly lit by the top and side monitors. The two areas were connected by a 7x7 cm gate in the center of the partition. Each animal was initially placed in a red plexiglass box positioned at the entrance of the light compartment and allowed to spontaneously enter the arena. Behavior was recorded for 10 minutes using infrared cameras and Bonsai software. The following parameters were extracted: total time spent in the dark area and the number of transitions between the dark and light areas. The mouse was considered in the dark zone when the body center coordinates were completely inside that area. The transitions were counted each time the body center coordinates cross the line between the two areas.

### Novel object Recognition Test

The test was divided into two sessions. On the first day, two identical objects were taped to the arena floor, the animal was gently placed in the arena and allowed to explore these objects for 10 minutes. As identical objects two green cubes (5 cm H x 5 cm W x 5 cm L) were used while the new object consisted of a little orange plastic watering can (7 cm H x 6 cm W x 3 cm L). On the second day, one of the objects was replaced with a new object and the mouse was put back into the arena for 10 min. The behavioral response was recorded with the bottom infrared camera and the was processed using DeepLabCut. The mouse was considered to be in proximity to an object when its body center coordinates were located within a circular area of 8 cm radius centered on that object. The discrimination index was defined as the difference in exploration time for the novel object (in sec) divided by the total exploration time (in sec)^73^.

### Social Interaction Test

The behavioral setup was divided with plexiglass walls into three equally spaced chambers and connected through 7x7 cm gates in the center of each wall. In the middle of the lateral chambers, two cages of 10 cm diameter made of transparent plexiglass grilles were placed. During the training phase of the experiment, mice were allowed to explore the arena for 10 minutes (training session). Immediately after, a novel conspecific was placed in one of the plexiglass cages, while the other remained empty. Again, the tested mouse was allowed to explore the setup freely for 10 minutes. We manually counted the time spent at each cage during the two phases of the experiment.

### Elevated Plus Maze Test

The day of the experiment the animal was moved in its home cage into the testing room 1 hour before the experiment for acclimatation. The elevated plus maze was made of dark polyvinyl plastic and consisted of a cross-shaped structure with two open, and two closed arms placed 50 cm above the floor, 55 cm long and 10 cm wide. The animal was gently placed in the central intersection area facing the open arm of the maze, and its activity was recorded with an infrared camera (Basler-60umPRO) placed on top of the apparatus for 10 min. The videos were manually scored to measure the time spent in the closed arms.

### Marble Burying Test

The test was performed in a standard rat cage filled with clean bedding material evenly distributed to a depth of approximately 5 cm. Twenty glass marbles (1.5–2 cm in diameter) were arranged on the bedding surface in a 4 × 5 grid pattern. Each mouse was gently placed in a corner of the cage, which was then covered with a filter top, and behavior was recorded from above for 30 min with an infrared camera (Basler ace acA1300-60gmNIR) using Bonsai software. At the end of the session, the mouse was carefully removed without disturbing the bedding, and the number of buried marbles was counted. A marble was considered buried when at least two-thirds of its surface was covered by bedding. Marbles were thoroughly cleaned after each trial, and a fresh cage with new bedding was used for each animal.

## IN VIVO NEUROPIXELS RECORDINGS

### Headpost surgery and craniotomy

Head-plate surgery was performed a week before the beginning of head-fixed experiments. Once the mouse was anesthetized with isofluorane (Iso-vet BRAND, NUMBER; 3% for induction, 1–3% during surgery) and placed on a stereotactic system (RWD), the skull was exposed, and Vetbond (3M, 7100139485) was applied to open skin and exposed muscle. A stainless-steel head plate (Neurotar, NTR000209-02) for headfixation was cemented to the skull (dental cement, Dental Leader, 12-909). During a separate surgery, cranial windows (∼0.5 to 1 mm2) were performed over the superior colliculus (SC) using a dental drill ( Marathon Italia, MA-005). The following coordinates were used as the center of craniotomy for the SC: AP: −3.7, ML: −0.5. Each mouse received a single injection of buprenorphine (Rymadil, 0.2 mg kg−1 subcutaneous injection) and antibiotics (Batyril; 1 ml per 100 ml) at the beginning of each surgery. The animals’ weight and health status were checked for 2 consecutive days.

### Headfixed set-up

The head-fixation apparatus consisted of the Mobile HomeCage system (HeadFix; Neurotar Ltd), in which head-fixed animals retain the possibility of freely moving on an ultralight 2D locomotion platform floating above an air-dispensing base. The floating platform consisted of a 35 cm diameter circle, with a magnetic locomotion-tracking system designed to record animal movements during the experimental session. Above the aluminum arm where the mouse was head-fixed, the 3D-printed Neuropixels probe holder (https://github.com/nerf-common/chronic-neuropixels-protocol) was in place connected to a micromanipulator (Sensapex, uMp-4). The probe was inserted at a precise speed of 0.002 mm/sec.

### Experimental procedure – Neuropixels Recordings

Starting from day 3 after the head-post surgery, each mouse was handled for 15 minutes by the experimenter for 3 consecutive days to reduce stress. The animal then underwent three sessions of habituation to the head-fixed setup. Each habituation session consisted of an hour in the head-fixed stage in front of two monitors with grey screens, while recording the animal movement. Following habituation, a craniotomy was performed above the SC as described above. Either later on the same day or on the next day, mice were placed in the head-fixed set-up and a Neuropixels probe 1.0 (imec) coated with a fluorescent dye (DiD, DiI; Thermofisher, D282, D307) was slowly inserted into the right or left cortex, SC and periaqueductal gray (PAG). The exposed brain was covered with saline solution (Sodium chloride 0.9%, Eurospital). Experiments started 20 minutes after inserting the probe. Visual stimuli were presented on a 24-inch LCD monitor (Samsung T35F, 1920x1080 pxl, 75 Hz refresh rate) with the lower part of the monitor placed 30 cm in front of the contralateral eye respective to the probe insertion side (covering 90° of azimuth and 70° of altitude ) and at an angle so that the distance between the eye of the mouse to the left corner, right corner and top of the monitor was similar. The setup was surrounded by black fabric to minimize surrounding light. Recordings were obtained from 384 electrodes at the tip of the probe, spanning a total of 3’840 µm in depth. Signals were acquired using the Neuropixels headstage and base station (imec) connected to a PXI acquisition chain composed of a PXI-1083 Chassis (NI, 787026-01) and Neuropixels PXI module (imec, PXIe_1000). High frequencies (>300 Hz) and low frequencies (<300 Hz) were acquired separately. Real time monitoring and data acquisition were performed with SpikeGLX software (V20230101-phase30, https://billkarsh.github.io/SpikeGLX). Experiment workflow was controlled using Bonsai, which generated synchronization pulses transmitted to an Arduino microcontroller (Arduino). The Arduino distributed the signals to the Neuropixels PXI module and to the Neurotar tracking system to ensure temporal alignment of neural and behavioral data. In addition, a photodetector precisely recorded the visual stimulus onset, ensuring accurate temporal synchronization across all systems. Each mouse underwent a maximum of two experimental sessions where the Neuropixels probe was inserted once into each hemisphere. Following the recordings, the mice were euthanized, the brain was extracted, fixed for at least 24h in 4% paraformaldehyde (Prodotti Gianni, P087.1). The brain was cut into 100-µm-thick slices using a vibratome (Campden Instrument, 5100 mz Vibrotome) and stained with DAPI (1:1000). The probe location was verified by confocal images of the fluorescent dye.

### Visual stimuli

Visual stimuli were programmed with Bonvision and presented on a grey background. During each session two stimuli were presented three times each in a randomized order: a black looming disk (from 2° to 50° visual angle in 250 ms; the disk stayed at full size for 250 ms and another black loom appeared after 500 ms) and a dimming disk (50° visual angle that changed from grey to black within 250 ms and remained for 250 ms and another dimming disk appeared after 500 ms). Each series of stimuli were presented on the monitor in front of the contralateral eye with respect to the insertion side of the Neuropixels probe.

### Spike sorting

SpikeInterface^74^ was used to pre-filter the high-frequency recording with high-pass filter set at 300 Hz. Eventually the data were spike-sorted with Kilosort 4^75^, followed by manual curation in Phy2. Units were classified as single units based on waveform-shape, refractory period violations assessed through autocorrelograms, and cluster isolation quality. Only good units were included in the analysis.

Anatomical borders between the retrosplenial cortex and the superficial superior colliculus (sSC), the sSC and deep superior colliculus (dSC), as well as between the dSC and periaqueductal gray (PAG), were defined by combining histological reconstruction with electrophysiological signatures as described previously^63^. The probe tract was reconstructed from fluorescent tracer labeling together with the known insertion depth. The upper borders of the sSC and dPAG were clearly identifiable in histological sections, allowing extraction of corresponding electrode positions.

### Neuropixels data processing

After spike sorting, spike times of each neuron were aligned with the visual stimulus onsets for further analysis. We then analyzed spontaneous and stimulus-evoked activity:

Burst analysis (Figure 4B-C): Similar to the MEA data, every burst produced by every single neuron in the periaqueductal gray and (presumably) retrosplenial cortex was extracted. A burst was defined as a sequence of at least 3 consecutive spikes with <8 ms inter-spike interval between each.

Correlation analyses (Figure 4D): Cross-correlations were computed identically to the analysis performed on MEA multi-unit data, except that no minimal firing rate threshold was applied.

Looming/dimming responses (Figure 5): For each cell the peri-stimulus spikes were collected, and a firing rate was calculated using 20 ms bins. Z-scores were computed based on the mean firing rate 1 s before stimulus onset. Only cells with a peak z-score of at least 2 during the first 2 s after stimulus onset were considered as ‘responding’. Mean and peak responses as well as area under the curve were calculated for these neurons only. For peak responses, the peak z-score during the first 2 s after stimulus onset was taken. The area under the curve was calculated during the first 3 s after stimulus onset.

### In vivo data model

Datasets from the in vivo Neuropixels recordings were analyzed via Hierarchical Bayesian regression^76^ to explicitly relate and disentangle single neuronal responses with respect to specific visual stimuli shown to the mice. There were four different visual stimuli, each presented twice in succession. Only results related looming and dimming were discussed in the results section; variables relating to white looming and sweeping were described in the supplementary information. For each neuron, the response activity (Y) to a visual stimulus was quantified as the difference in the total number of spikes within the time windows just before and during a stimulus presentation (0.5s each). The binary design matrix X described all possible stimulations considered. It consisted of eight columns: two for each stimulus (of a total of four), with one referring to the type of stimulus shown and one marking the second time each stimulus was shown in succession. To each column t was fitted a coefficient Bt and the linear intercept was set to zero as a ‘null response’. We assumed normal dispersions

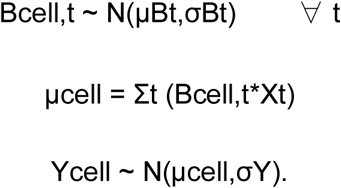

where μBt and σBt were the mean and standard deviation for the distribution of coefficients Bt across neurons, and σY was the observation noise. For numerical stability we used a non-centered parametrization so that Bcell,t = μBt + τ*σBt, with τ∼N(0,1). Three models with different hierearchical structures were considered. In the “single prior” model, a single global prior was assigned for each of the eight coefficients:

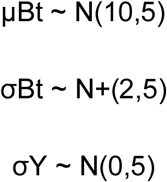

In the “genotype prior” model, μBt and σBt were separated into two distinct distributions, one for WT cells and one for KO cells:

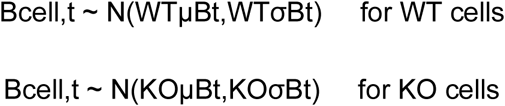

In the “separate mice” model, μBt was separated between a mean component common to all cells, and a component specific for each mouse:

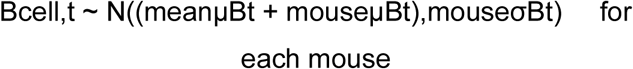

The posteriors were sampled in 8 independent chains of 2000 iterations (of which the first 1000 were discarded for tuning), with a target acceptance rate of 0.8. We qualified the model fits using Expected Log-likelihood Probability Density (ELPD), which is a measure of the model predictive power over unobserved data. ELPD of the different models were compared via Pareto smoothed importance sampling leave-one-out cross-validation^77,78^. These comparisons were performed using arviz 0.21.0^79^. Statistical significance of the difference between posterior means was determined via “probability of direction”^80^ with the threshold α = 0.05 (ns; * p < 0.05; ** p < 0.01; *** p < 0.001).

Additional information on the mice movement during each trial (average instantaneous acceleration) only resulted in minimal ELPD increases, so it was not included in the main models (Figure S7). Diagnostics information for μB and σB can be found in Table S2.

## QUANTIFICATION AND STATISTICAL ANALYSIS

Statistical analyses of molecular *in vitro* data were performed in GraphPad Prism (v8.0.2), and results were based on two-tailed t-test. *In vitro* electrophysiological evaluations were performed in MATLAB (MathWorks), and specific data analysis is described above. *In vivo* data analysis was based on estimation statistics using the DABEST package ^81^ and Wilcoxon Ranksum Test. To compare escape probability across genotypes, a generalized linear the mixed-effects model with a binomial distribution and a logit link function was fit accounting for mouse identity.

Significance threshold was α = 0.05 (ns; * p < 0.05; ** p < 0.01; *** p < 0.001; **** p < 0.0001).

## MATHEMATICAL MODEL

To explore putative subcellular mechanisms, underlying spontaneous bursting observed in the in vitro experiments, we computer simulated an excitatory-inhibitory (E-I) firing-rate mathematical model (Wilson & Cowan, 1972), following previous studies (Moskalyuk et al., 2020). The state of such a network is described by the mean firing rates of the excitatory (*r_E_*) and inhibitory (*r_I_*) populations. We also introduced a slow adaptation variable (*a*), representing activity-dependent negative feedback mechanisms (e.g., intrinsic slow *K*^+^ ion currents such as *I_K_*_,*Ca*_, or short-term synaptic depression), affecting the excitatory population only. The system was described by three coupled differential equations:

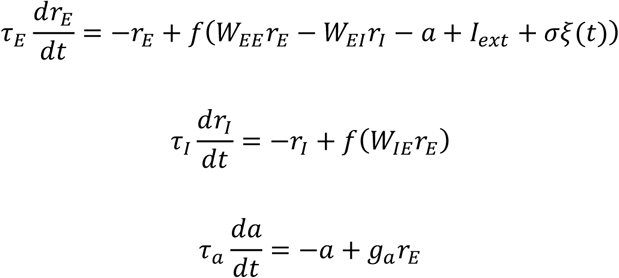

where *τ_E_* and *τ_I_* are the fast membrane time constants for the excitatory and inhibitory populations, respectively, and *τ_a_* ≫ *τ_E_*, *τ_I_* is the time constant of the slow adaptation variable. The parameters *W_XY_* denote the average synaptic efficacy from population *Y* to population *X*. *I_ext_* represents a stationary baseline drive, due to background feedforward afferents not explicitly modelled, also capturing spontaneous synaptic release or persistent sodium currents.

While recurrent glutamatergic connections *W_EE_* act as positive feedback and primarily modifies the frequency burst occurrence, *g_a_* dictates the rate of activity-dependent fatigue, effectively terminating each burst. The firing rates are finally bounded by an activation function *f*(*x*), approximating the input-output properties of each neuron (i.e. the f-I curves) as a sigmoid:

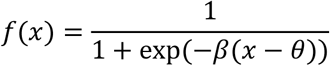

where *β* defines the gain and *θ* is the firing threshold. To account for fluctuations associated to finite-size networks and random synaptic release, we introduced a Gaussian noise term *ξ*(*t*) to the external drive, with zero mean and unit variance, scaled by the fluctuation amplitude *σ*.

## SUPPLEMENTARY MATERIAL

**Figure S1.**
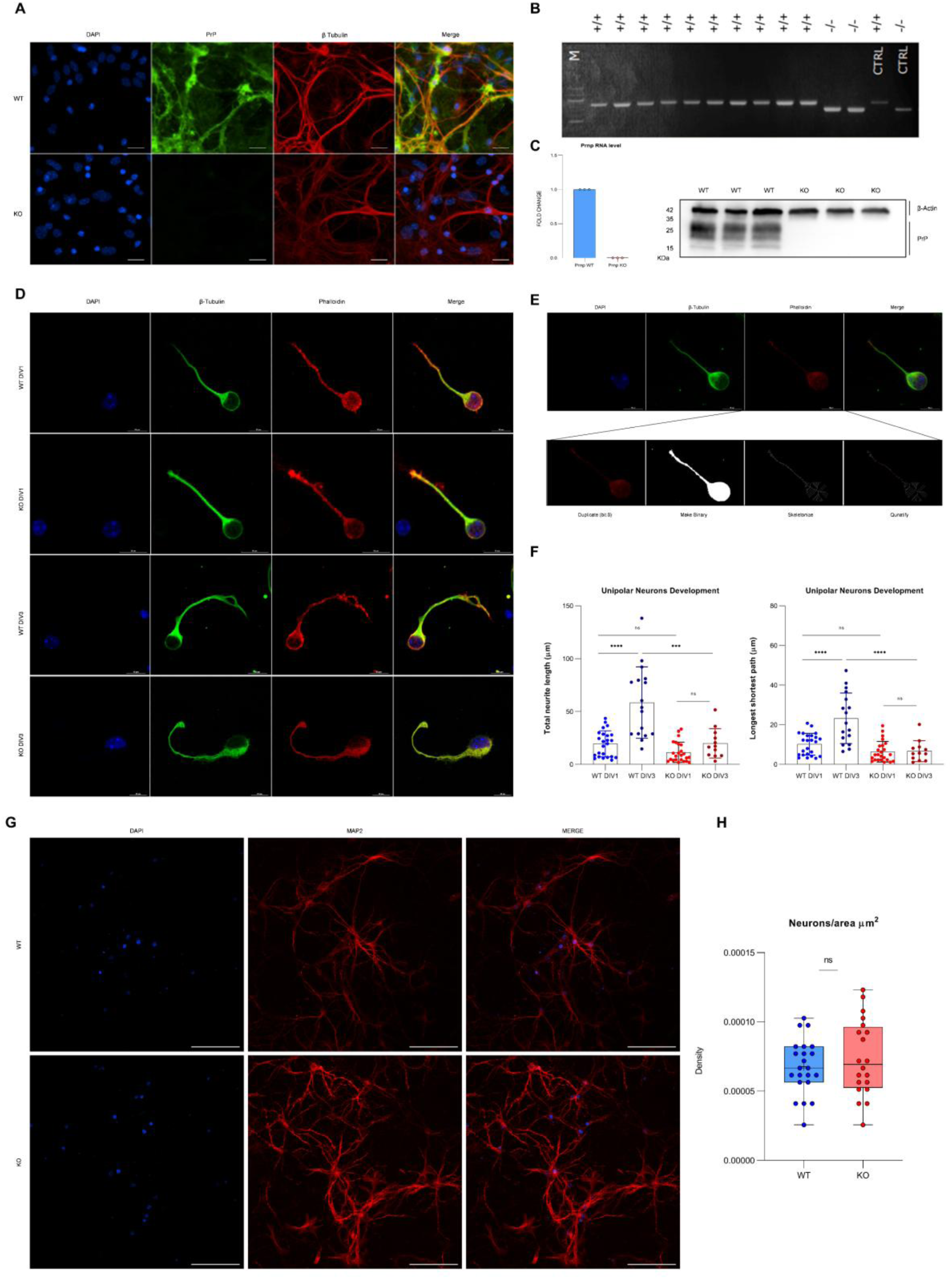
FVB model characterization. (A) Representative immunostaining of primary neuronal cultures showing absence of PrP (green) in KO compared with WT; nuclei (DAPI, blue) and β-tubulin (red); merged images shown (scale bar 100 µm). (B) PCR genotyping confirming KO and WT alleles, including internal controls. (C) Loss of Prnp expression confirmed at the RNA level by RT–qPCR (left) and at the protein level by Western blot (right) in three independent biological replicates (β-actin loading control shown). (D) Representative DIV1 and DIV3 unipolar neurons stained with β-tubulin and phalloidin to visualize neurites and F-actin, respectively; merged images shown (scale bars 10 µm). (E) ImageJ-based skeletonization workflow used for neurite quantification (original signal, binarization, skeletonization, and graph/longest-path extraction). (F) Quantification of total neurite length and longest shortest path in WT and KO unipolar neurons at DIV1 and DIV3. (G) Representative fields showing neuronal network organization (MAP2, red) and nuclei (DAPI, blue) in WT and KO cultures (scale bars 100 µm) at DIV23. (H) Neuronal density (neurons/area, µm²) in WT and KO cultures. Statistics: Two-tailed t test; * p < 0.05, ** p < 0.01, *** p < 0.001, **** p < 0.0001; n.s., not significant. Sample sizes: WT n = 7, KO n = 6 independent cultures.

**Figure S2.**
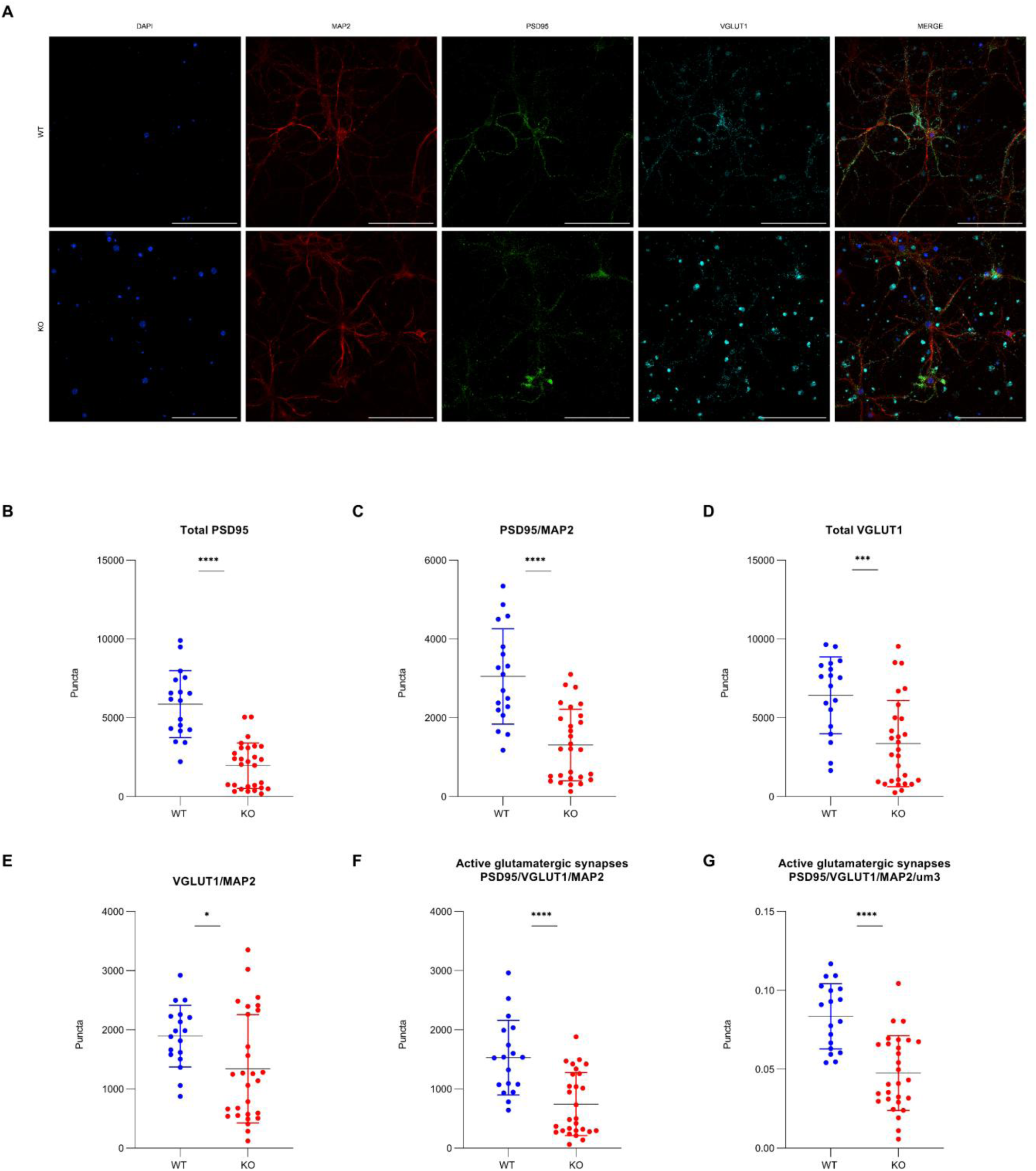
Immunostaining of PSD95 and VGLUT1. (A) Representative images of WT and KO primary neuronal cultures stained for nuclei (DAPI, blue), dendrites/neurites (MAP2, red), the postsynaptic marker PSD95 (green), and the presynaptic glutamatergic marker VGLUT1 (cyan), with merged channels shown (scale bars 100 µm). (B) Total PSD95 puncta. (C) PSD95 puncta normalized to MAP2 (PSD95/MAP2). (D) Total VGLUT1 puncta. (E) VGLUT1 puncta normalized to MAP2 (VGLUT1/MAP2). (F) Active glutamatergic synapses quantified as PSD95/VGLUT1 colocalized puncta within MAP2-positive neurites (PSD95/VGLUT1/MAP2). (G) Active glutamatergic synapse density (PSD95/VGLUT1/MAP2 per µm³). Each dot represents an independent field/ROI; bars indicate mean ± SD. * p < 0.05, ** p < 0.01, *** p < 0.001, **** p < 0.0001; ns, not significant.

**Figure S3.**
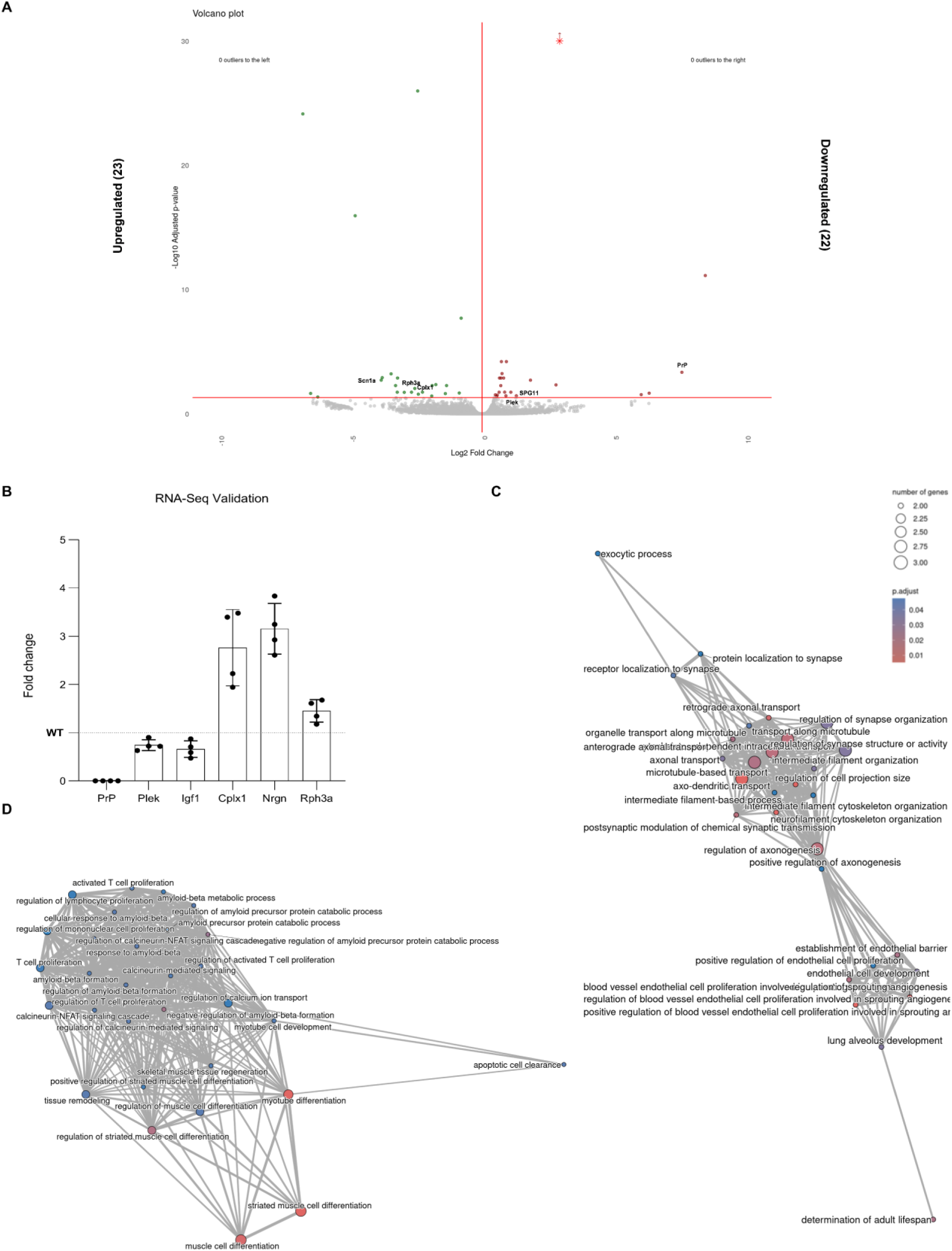
RNA-Seq of DIV23 FVB primary cultures (KO vs WT). (A) Volcano plot of differential gene expression (x-axis: log2 fold change KO/WT; y-axis: −log10 FDR). Genes passing the significance and fold-change thresholds (red guide lines) are highlighted; 23 genes are significantly upregulated and 22 significantly downregulated in KO cultures. (B) RT-qPCR validation of selected differentially expressed genes: the three most downregulated (PrP, Plek, Igf1) and the three most upregulated (Cplx1, Nrgn, Rph3a) from the RNA-seq analysis. Bars show fold change relative to WT (dotted line), with individual biological replicates overlaid (mean ± variability as shown). (C) Gene Ontology (GO) enrichment network for upregulated genes. Nodes represent enriched GO Biological Process terms; node size reflects the number of genes in the term and node color indicates adjusted enrichment significance (FDR). Edges denote term overlap/similarity. (D) GO enrichment network for downregulated genes (visualization as in C).

**Table S1.**
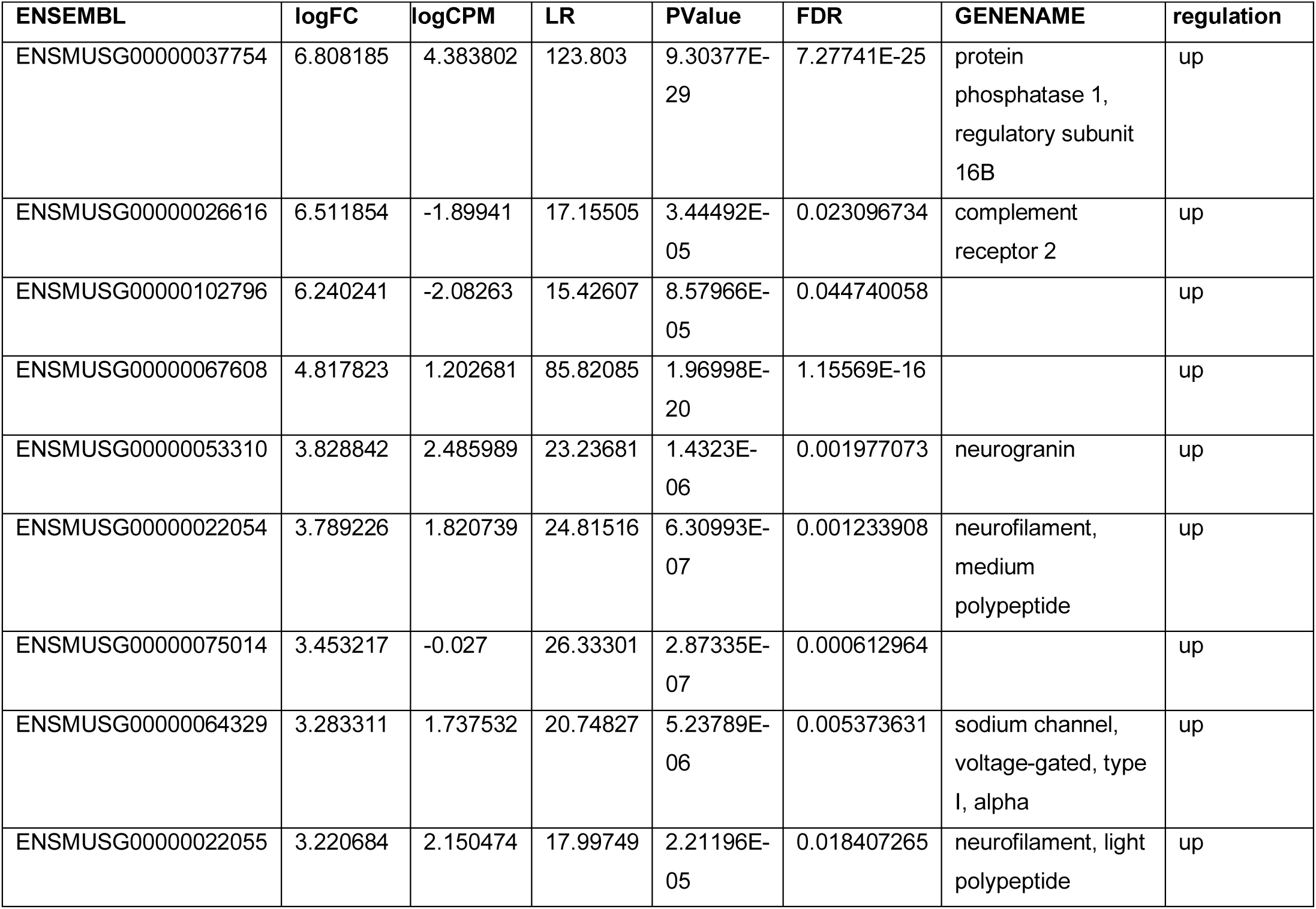
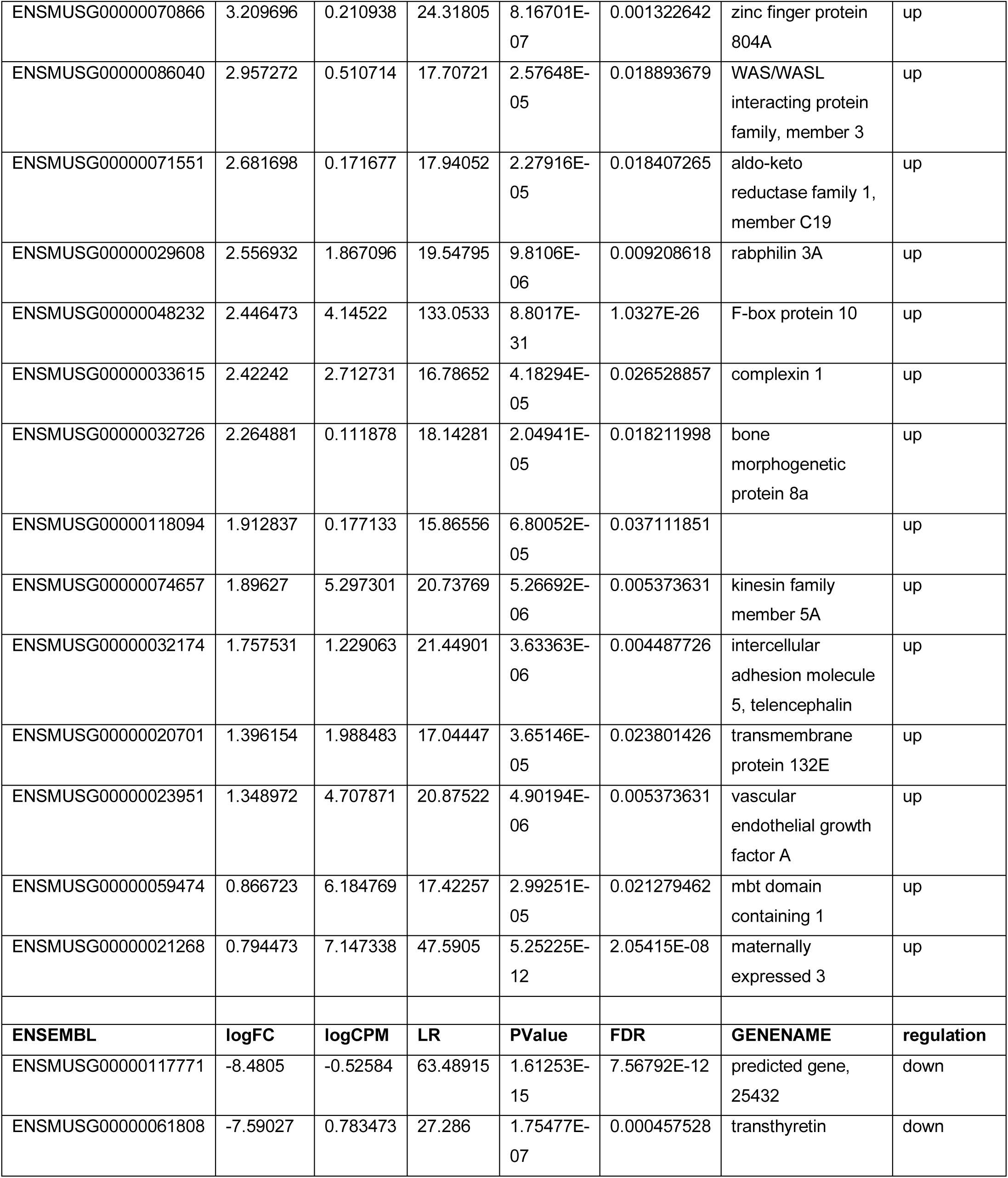
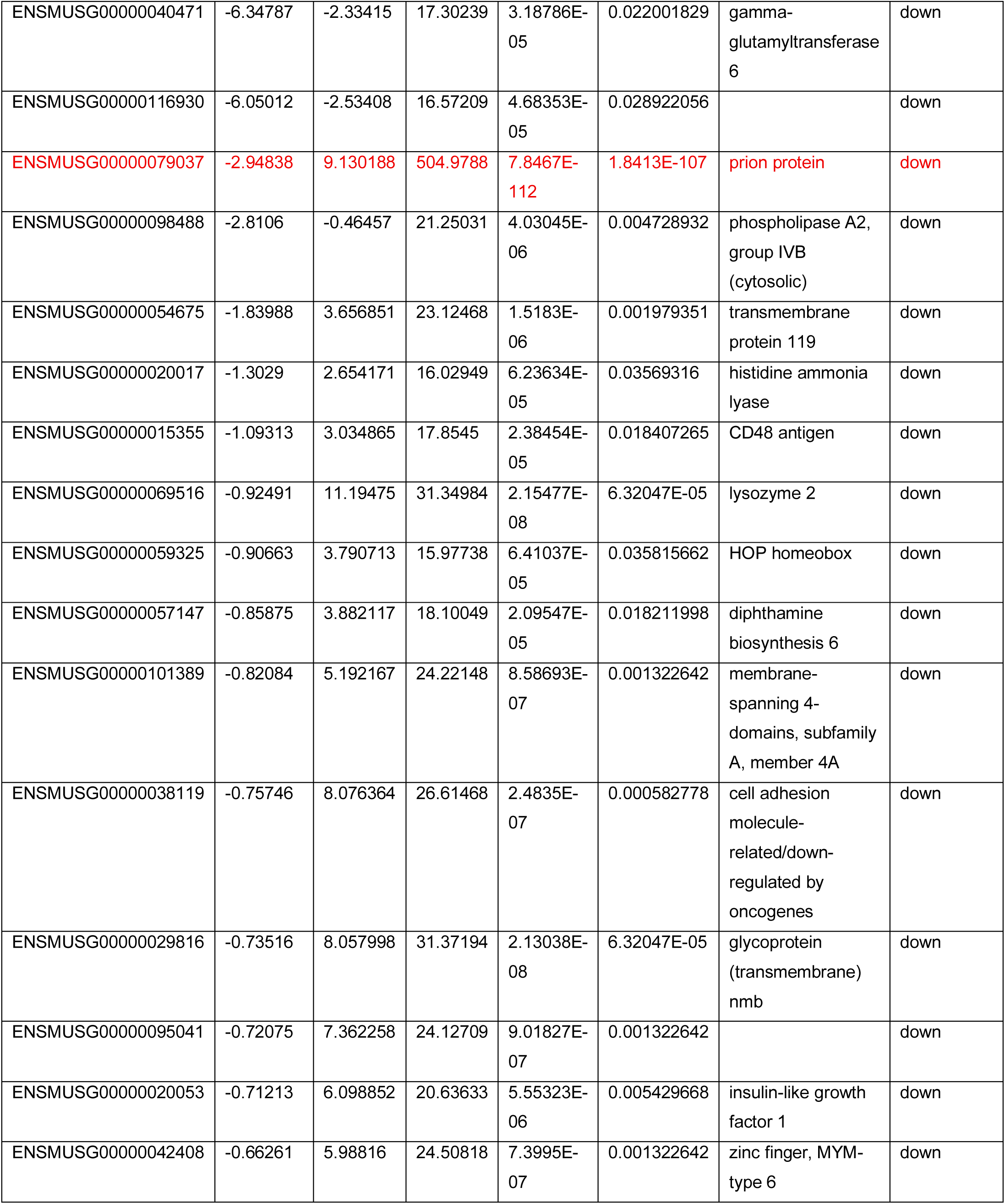
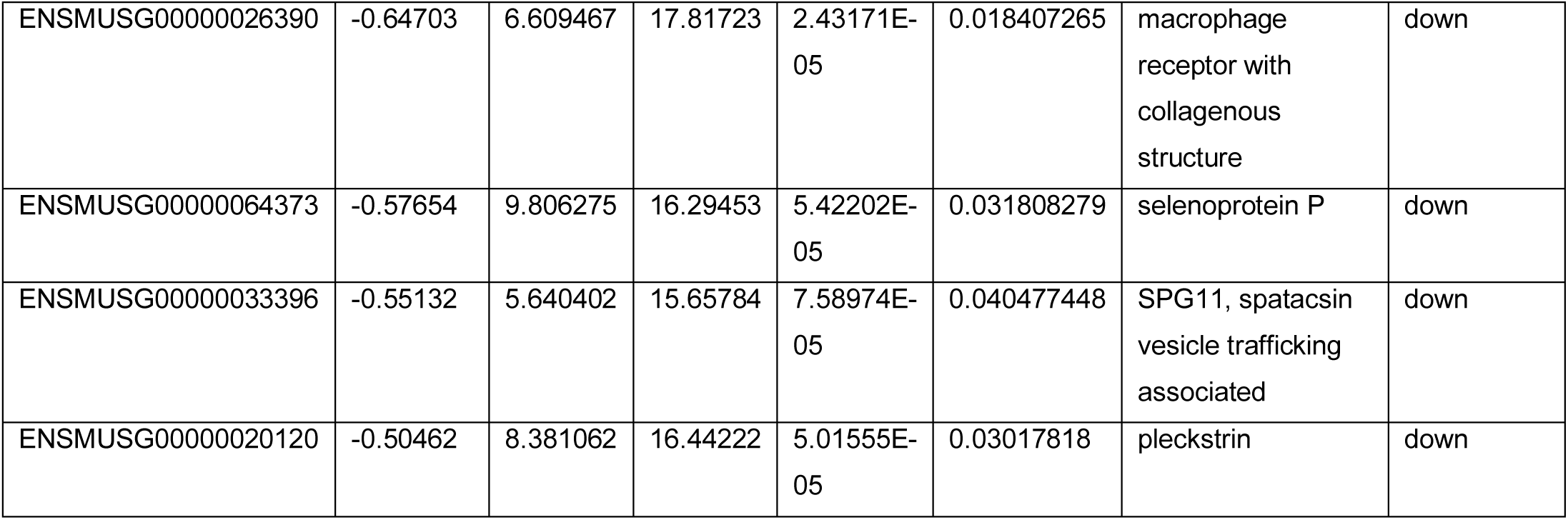
Differentially expressed genes in DIV23 FVB primary cultures (KO vs WT), shown as upregulated (top) and downregulated (bottom). Columns (in order): Ensembl ID; logFC (log2 fold change); logCPM (log counts per million); LR (likelihood ratio test statistic); P value; FDR (multiple-testing adjusted P value); gene name; and regulation direction. Prnp (prion protein) is highlighted in red.

**Figure S4.**
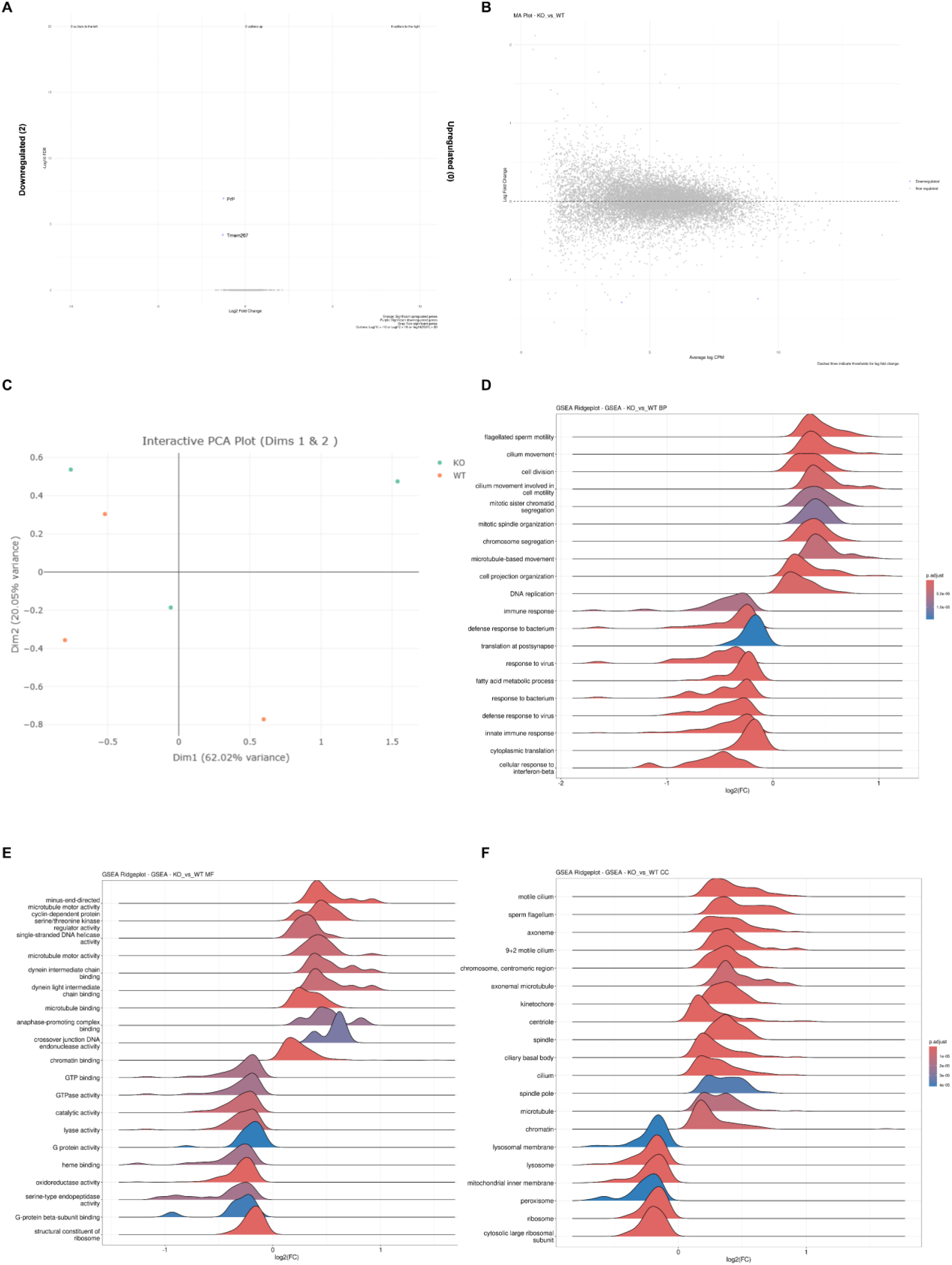
RNA-Seq of DIV23 ZH3 primary cultures (KO vs WT). (A) Volcano plot revealing 0 upregulated and 2 downregulated genes in KO cultures. (B) Bland-Altman Plot showing log2 fold change (KO/WT) as a function of average expression (logCPM). (C) PCA of normalized expression values shows no clear separation between KO and WT samples (D-F) GSEA using ranked gene lists reveals enriched gene sets in Biological Process (D), Molecular Function (E), and Cellular Component (F) categories.

**Figure S5:**
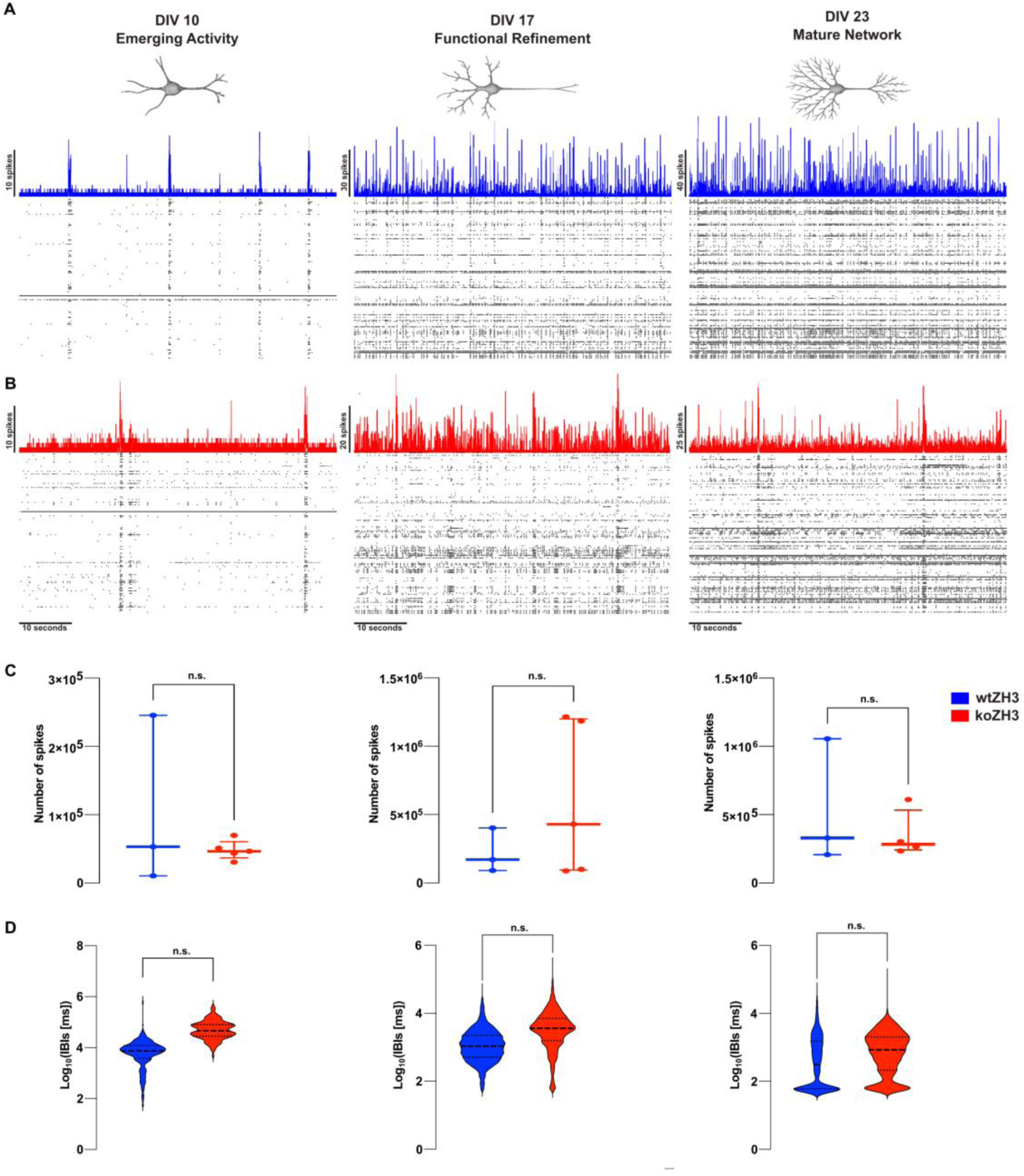
Network-burst occurrence across development in PrP-null ZH3 cortical cultures. **(A–B)** Representative multielectrode-array (MEA) recordings from neocortical cultures at DIV10, DIV17, and DIV23 showing spike rasters (black) with the corresponding spike-time histograms (STHs; 5-ms bins; wtZH3, blue in A; koZH3, red in B). Scale bars as indicated. **(C)** Total spike count per MEA during 30-min spontaneous recordings at each developmental stage. Each dot represents one MEA; horizontal lines indicate median with interquartile range (IQR). DIV10: unpaired t test with Welch’s correction, two-tailed, p = 0.6635 (wtZH3 n = 3 MEAs, koZH3 n = 5 MEAs). DIV17: unpaired t test with Welch’s correction, two-tailed, p = 0.0774 (wtZH3 n = 3, koZH3 n = 5). DIV23: unpaired t test with Welch’s correction, two-tailed, p = 0.5515 (wtZH3 n = 3, koZH3 n = 5). **(D)** Inter-burst interval (IBI) distributions for network-wide bursts at DIV10, DIV17, and DIV23. Violin plots display log10-transformed IBI values (ms) for visualization only; statistics were performed on raw, untransformed IBI values. DIV10: clustered Wilcoxon, adjusted p = 0.5893 (wtZH3 n = 3 MEAs, 392 IBIs; koZH3 n = 5 MEAs, 120 IBIs; median of MEA medians = 14,400 vs 68,053 ms; r_rb = −0.405). DIV17: clustered Wilcoxon, adjusted p = 0.5893 (wtZH3 n = 3 MEAs, 2525 IBIs; koZH3 n = 5 MEAs, 1280 IBIs; median of MEA medians = 1453 vs 3468 ms; r_rb = −0.362). DIV23: clustered Wilcoxon, adjusted p = 0.6000 (wtZH3 n = 3 MEAs, 2417 IBIs; koZH3 n = 3 MEAs, 1669 IBIs; median of MEA medians = 1545 vs 1185 ms; r_rb = −0.330). Effect size is reported as the cluster-weighted rank-biserial correlation (r_rb). Data from 1 independent culture preparation. *p ≤ 0.05; **p ≤ 0.01; ***p ≤ 0.001; ****p ≤ 0.0001; n.s., not significant.

**Figure S6:**
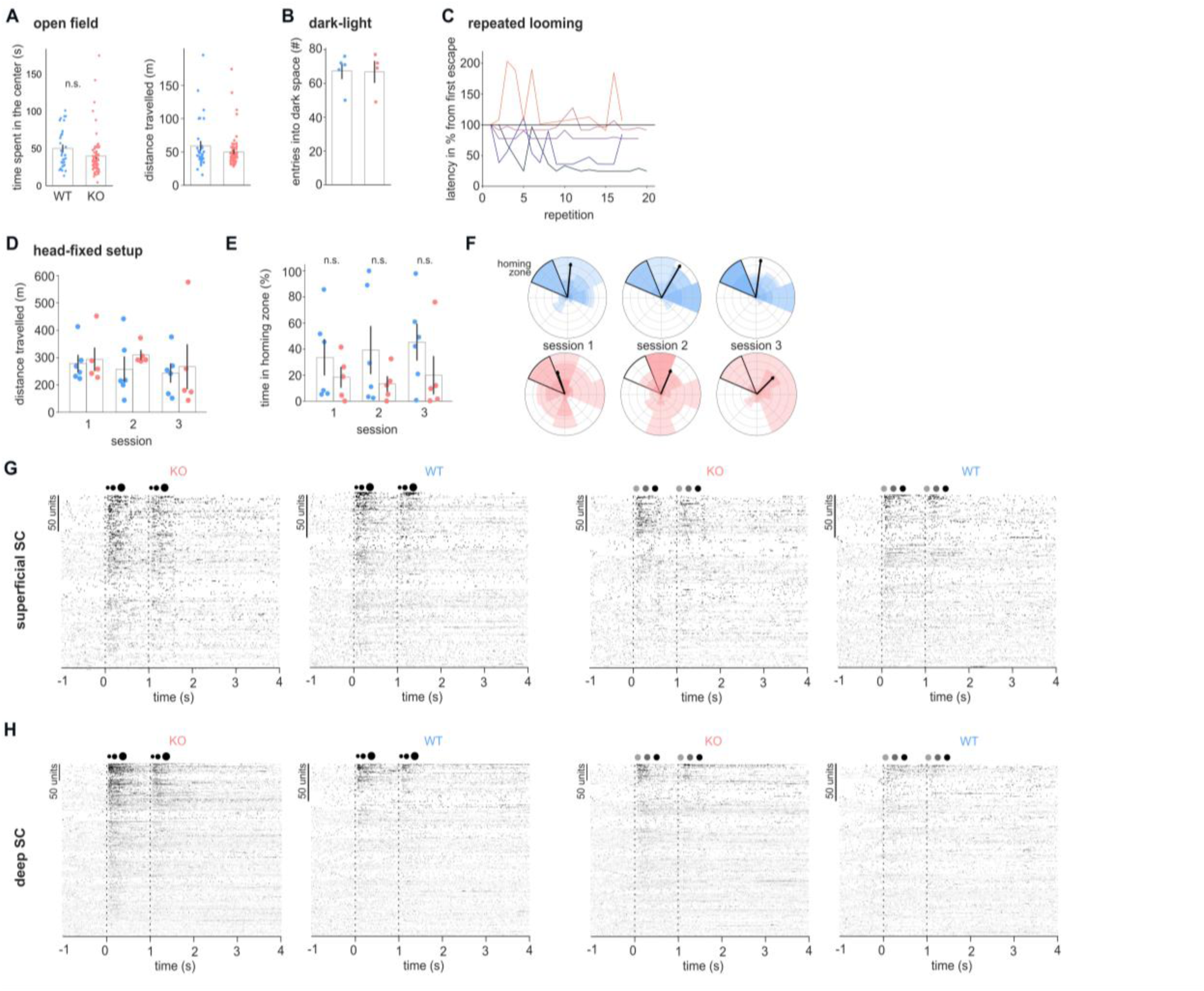
Additional information about in vivo experiments. (A) Open field assay. Time spent in the center of the arena and distance travelled for the same animals as in Figure 4. (B) Dark-light chamber. Number of entries into the dark compartment of the setup for the same animals as in Figure 4. (C) Latency to escape for repeated looming stimuli related to Figure 4L. Each line indicates the latency for a single animal to each of the 20 looming repetitions in percent relative to the escape latency for the very first loom. (D) Head-fixed setup. Distance travelled during each session for the same animals as in Figure 4M-N. (E) Raster plots of all recorded neurons in the superficial superior colliculus for looming (left) and dimming stimuli (right). Neurons were sorted by their peak response. Related to Figure 5E-F. (F) As in E but for neurons of the deep layers of the superior colliculus.

**Figure S7.**
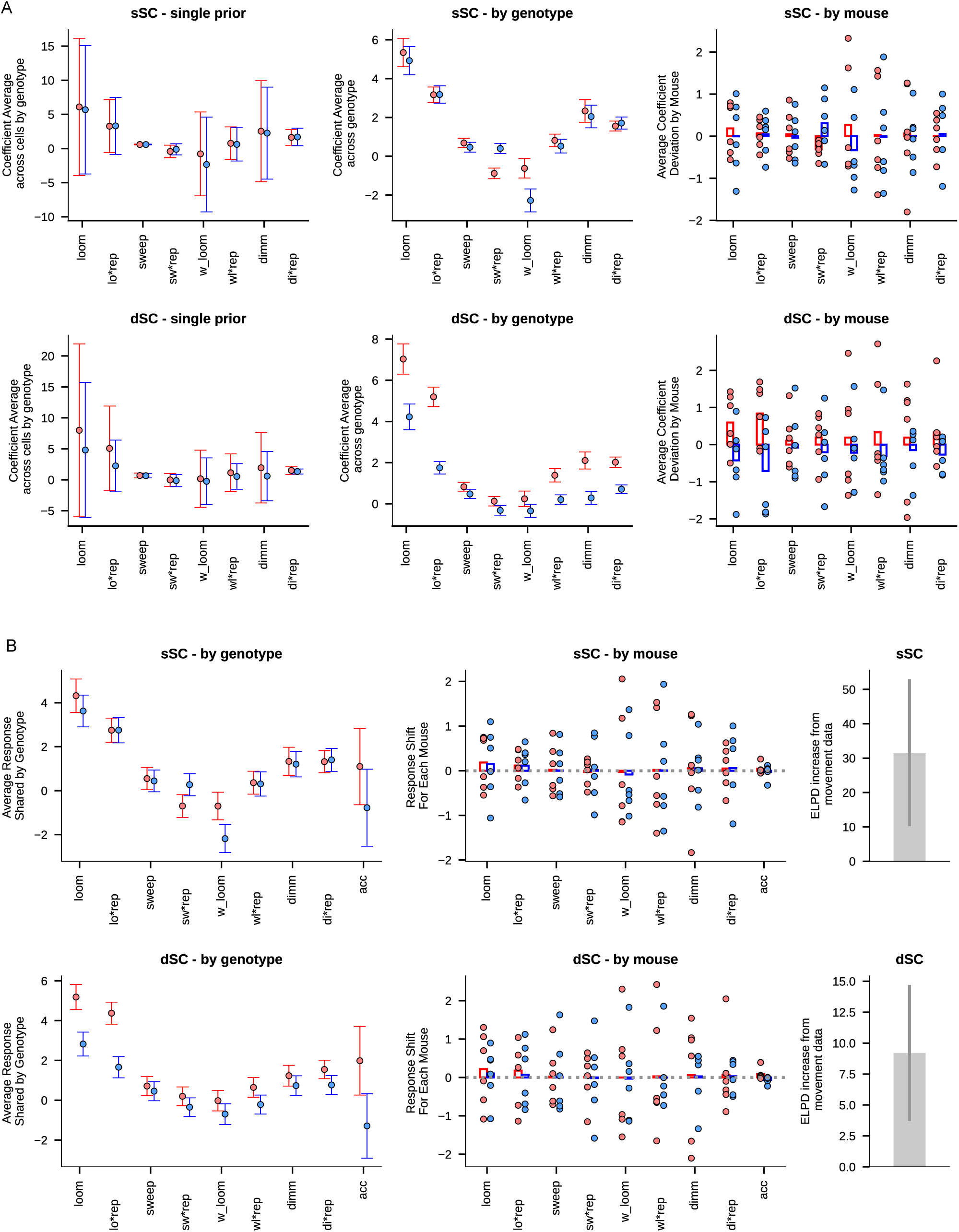
Full results of the in vivo models. (A) Same data from figure 5 G and H showing all coefficients including extra stimuli sweep and white looms. (B) Equivalent models to (A) that include an extra variable average instantaneous acceleration of mouse movement during each stimulus. A discreet increase of ELPD was observed, which is compatible with the notion of locomotion information representation in the SC.

**Table S2.**
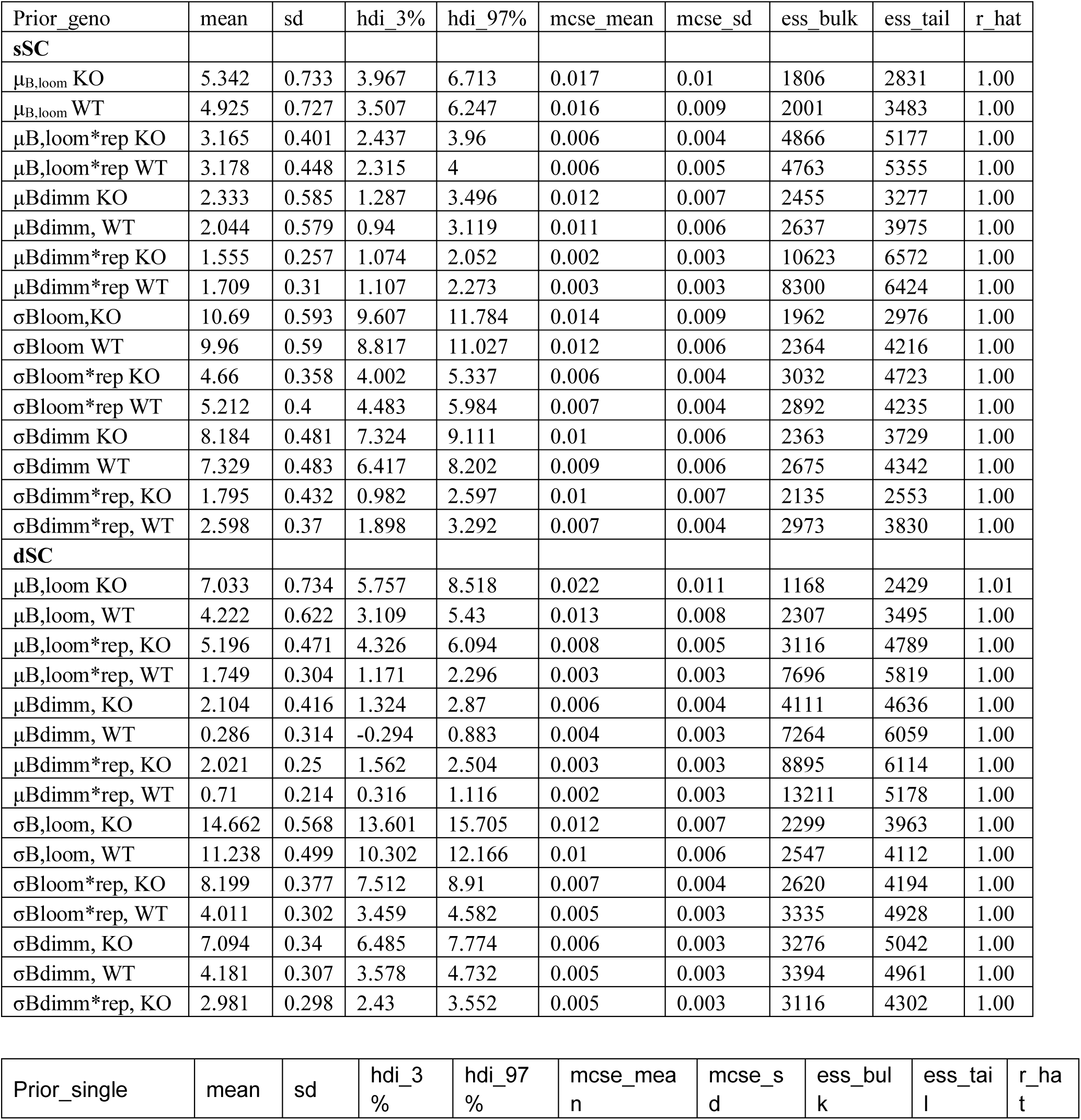
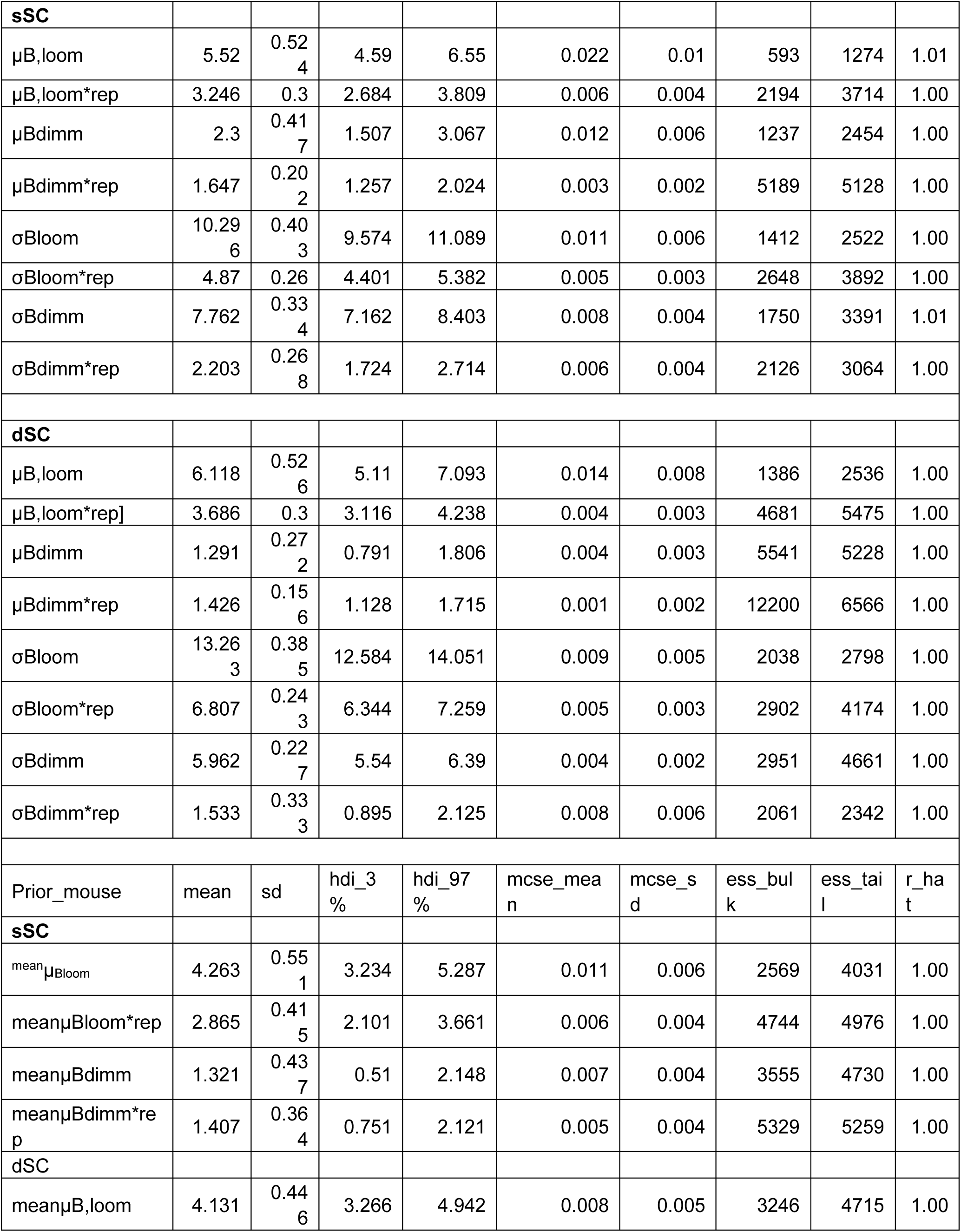
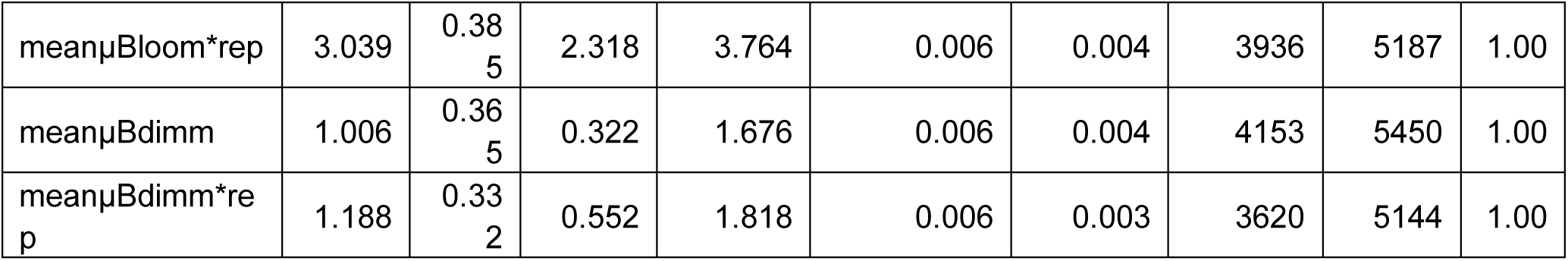
Diagnostics of the fitted posterior distributions from the three in vivo models. Statistics using prior_geno (top), prior_single, and prior_mouse (bottom), each for sSC and dSC. Variables shown correspond to average (µ) and standard deviation (σ) of coefficients for looming and dimming stimuli in each category. The following values were calculated across samlples for each variable: mean, standard deviation, highest density intervals (HDI 3 and 97%), Monte Carlo Standard Error Mean and standard deviations, and convergence effective sample sizes (ess) and r_hat.

**Figure S8.**
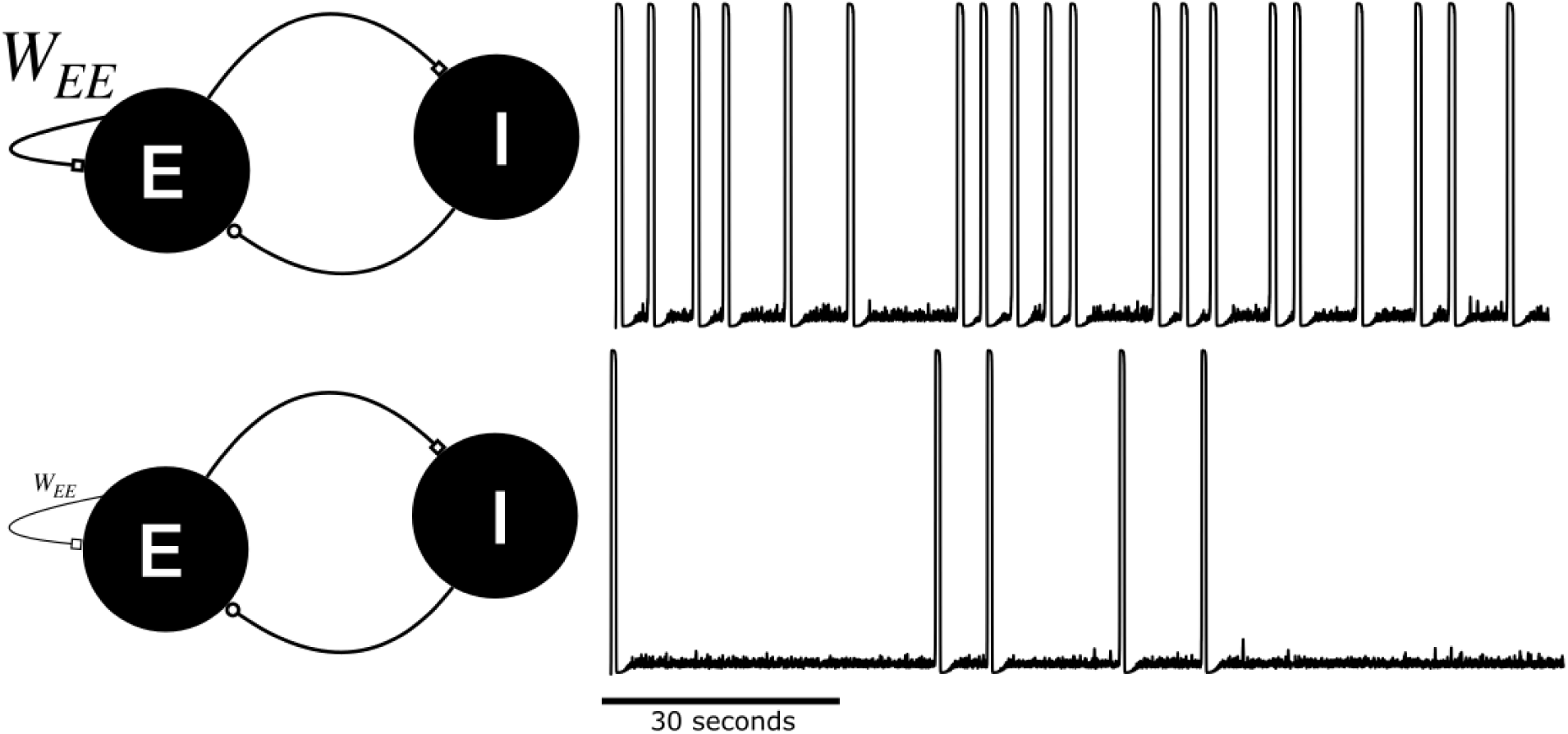
Spontaneous network bursting reflects recurrent excitatory connections numerosity or efficacy. In a simplified mathematical model of excitatory-inhibitory (E-I) network, the population firing rate displays spontaneous bursting. The rate of episodic bursting reflects the recurrent excitatory synaptic drive W_EE_, which acts as positive feedback and transiently destabilizes the network. Identical networks simulated with a strong **(top)** or weak **(bottom)** drive displays shorter or longer inter-burst intervals, respectively.

**Table S3.**
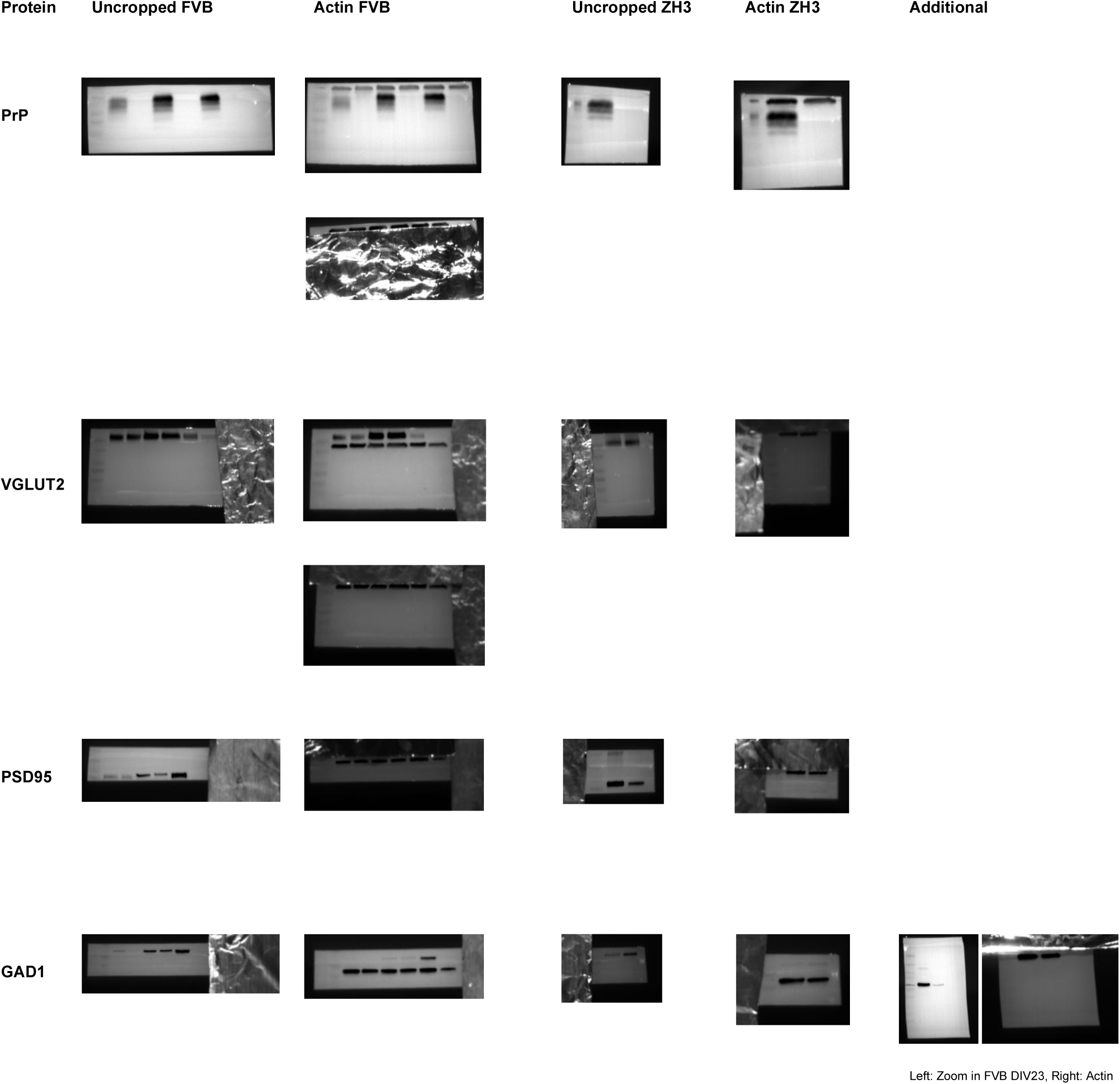
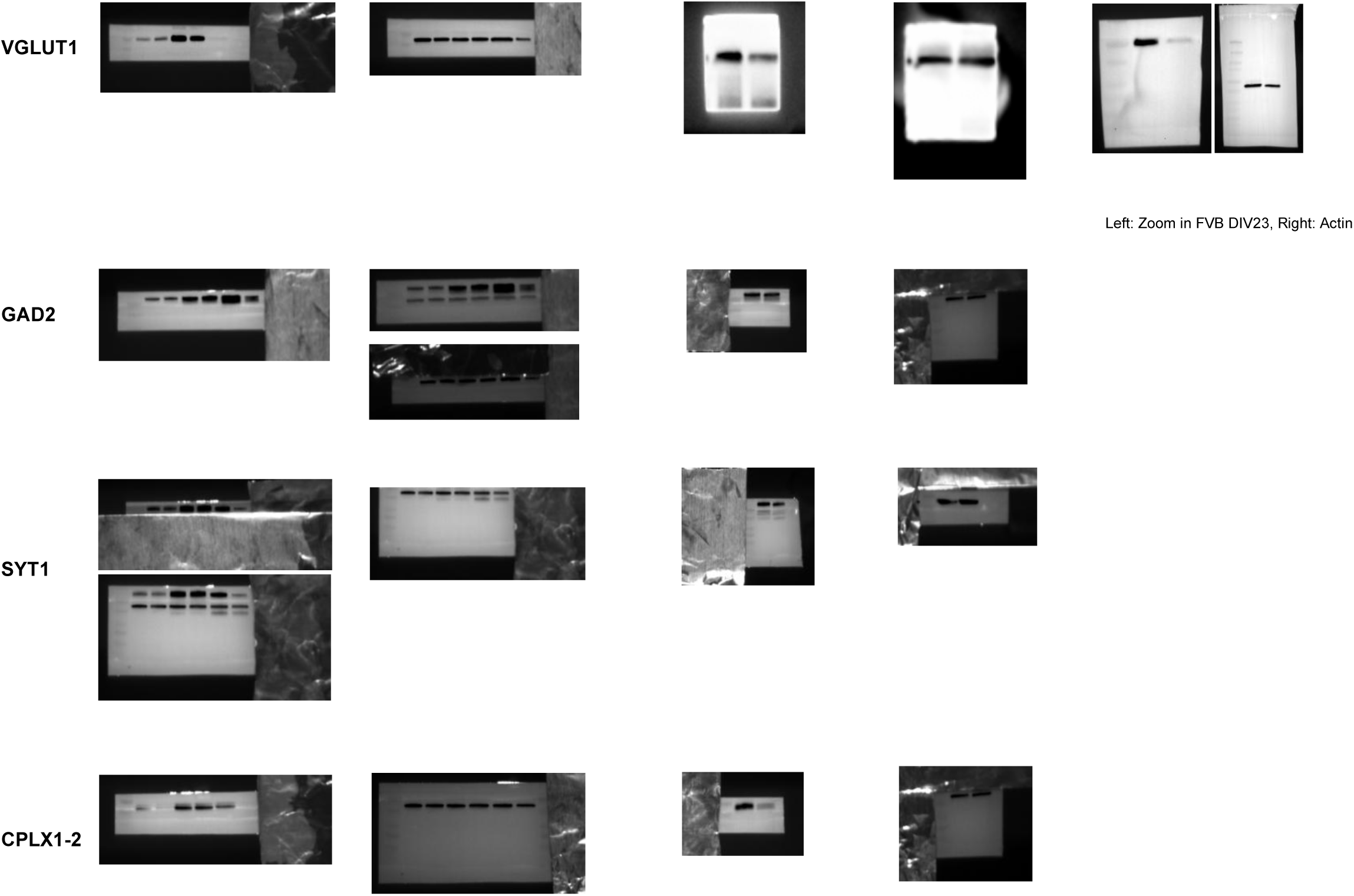
Uncropped blots. Loading order FVB: WT_DIV10 – KO_DIV10 – WT_DIV17 – KO_DIV17 – WT_DIV23 – KO_DIV23; Loading order Zoom in FVB: WT_DIV23 – KO_DIV23; Loading order ZH3: WT – KO

